# Rare GPR37L1 variants reveal potential roles in anxiety and migraine disorders

**DOI:** 10.1101/2023.07.05.547546

**Authors:** Gerda E. Breitwieser, Andrea Cippitelli, Yingcai Wang, Oliver Pelletier, Ridge Dershem, Jianning Wei, Lawrence Toll, Bianca Fakhoury, Gloria Brunori, Raghu Metpally, David J. Carey, the Regeneron Genetics Center, Janet Robishaw

## Abstract

GPR37L1 is an orphan receptor that couples through heterotrimeric G-proteins to regulate physiological functions. Since its role in humans is not fully defined, we used an unbiased computational approach to assess the clinical significance of rare *GPR37L1* genetic variants found among 51,289 whole exome sequences from the DiscovEHR cohort. Briefly, rare *GPR37L1* coding variants were binned according to predicted pathogenicity, and analyzed by Sequence Kernel Association testing to reveal significant associations with disease diagnostic codes for epilepsy and migraine, among others. Since associations do not prove causality, rare *GPR37L1* variants were then functionally analyzed in SK-N-MC cells to evaluate potential signaling differences and pathogenicity. Notably, receptor variants exhibited varying abilities to reduce cAMP levels, activate MAPK signaling, and/or upregulate receptor expression in response to the agonist prosaptide (TX14(A)), as compared to the wild-type receptor. In addition to signaling changes, knockout of GPR37L1 or expression of certain rare variants altered cellular cholesterol levels, which were also acutely regulated by administration of the agonist TX14(A) via activation of the MAPK pathway. Finally, to simulate the impact of rare nonsense variants found in the large patient cohort, a knockout (KO) mouse line lacking *Gpr37L1* was generated, revealing loss of this receptor produced sex-specific changes implicated in migraine-related disorders. Collectively, these observations define the existence of rare GPR37L1 variants in the human population that are associated with neuropsychiatric conditions and identify the underlying signaling changes that are implicated in the *in vivo* actions of this receptor in pathological processes leading to anxiety and migraine.

**SIGNIFICANCE STATEMENT:** G-protein coupled receptors (GPCRs) represent a diverse group of membrane receptors that contribute to a wide range of diseases and serve as effective drug targets. However, a number of these receptors have no identified ligands or functions, i.e., orphan receptors. Over the past decade, advances have been made, but there is a need for identifying new strategies to reveal their roles in health and disease. Our results highlight the utility of rare variant analyses of orphan receptors for identifying human disease associations, coupled with functional analyses in relevant cellular and animal systems, to ultimately reveal their roles as novel drug targets for treatment of neurological disorders that lack wide-spread efficacy.

## Introduction

Migraine is a common neurovascular disorder with significant personal, societal and medical costs that affects 12-15% of adults, and has a strong sex bias toward females [1,2]. Migraine frequency, and the associated symptoms (e.g., severe headache, sensitivity to sensory inputs, nausea, and vomiting), vary among individuals, and can occur multiple times per week to several times per year. Dissection of the neurobiology of migraine has focused on increases in neuronal or cortical hyperexcitability [3,4] and/or changes in the trigeminal-vascular system of the dura [4,5]. While mechanism(s) have been defined once the attack begins, the causal basis for susceptibility to migraine remains poorly understood.

Genetic differences account for up to 60% of migraine occurrence [3,6–9]. Dissection of the underlying genetics has proceeded along two independent paths. At one end of the spectrum, <u>G</u>enome-<u>W</u>ide <u>A</u>ssociation <u>S</u>tudies (GWAS) and subsequent meta-analyses of migraine have identified common markers (<u>S</u>ingle <u>N</u>ucleotide <u>P</u>olymorphisms, SNPs) in or near genes associated with glutamate signaling (*MTDH/AEG-1, LRP1, MEF2D*), synaptic development (*ASTN2, FHL5*), endothelial cell and blood vessel wall function (*PHACTR1, MMP16, AJAP1, TSPAN2, TGFBR2*), pain pathways (*TRPM8*), or metabolism (*PRDM16, C7orf10*) [8]. Since these SNPs are commonly found in intronic or intergenic regions, assigning GWAS loci to adjacent genes and defining pathologic mechanisms has been challenging. At the other end of the spectrum, familial studies have identified rare mutations in specific genes (*NOTCH3, TREX1, CACNA1A, ATP1A2, SCN1A*) associated with monogenic hemiplegic migraine [3,4,9]. Since the rare variants are more often present in coding regions, an understanding how these mutations directly affect protein function and contribute to migraine has begun to emerge from genetic mouse models [10]. Although these strategies provide important mechanistic insights, the cumulative impact of the known genes accounts for a small fraction of estimated migraine heritability.

To address the “missing” heritability, recent attention focuses on the possible contribution of genes carrying rare variants that are not captured by GWAS genotyping assays and exert moderate effects not easily detected by classical linkage analysis in familial studies [11]. As a test of this concept, we selected a subset of 85 G Protein-Coupled Receptors (GPCRs) [12] whose endogenous ligands, biological functions and disease contributions have not been fully elucidated, i.e., orphan or recently adopted GPCRs [13,14]. We examined whole-exome sequences (WES) from 51,289 individuals, predicted likely pathogenic variants using bioinformatics algorithms, and then binned “likely pathogenic” plus “pathogenic” variants for each of the 85 GPCR genes in the study. Subsequently, Sequence Kernel Association Testing (SKAT) was conducted to determine the genes that significantly associated with the International Classification of Disease Diagnostic (ICD) codes found in the electronic health records (EHRs) of these individuals [12]. Among the top hits, the *GPR37L1 (G Protein-Coupled Receptor 37-Like 1)* gene, coding for a recently deorphanized GPCR, was significantly associated with generalized epilepsy [12]. Providing proof-of-concept for this unbiased approach, a point mutation in *GPR37L1* has previously been found in a consanguineous family with a progressive form of myoclonus epilepsy [15]. Attesting to the discovery potential of this approach, we report here that variation within the *GPR37L1* gene was also significantly associated with migraine. The encoded GPR37L1 receptor is highly expressed in astrocytes and microglia [16–18]. Although it has been postulated to contribute to neuroprotection [16–18], the cellular pathways have been poorly defined. In this paper, functional characterization of migraine-associated missense variants in GPR37L1 was conducted revealing agonist-induced alterations in expression, signal amplitude, and pathway bias. In addition to signaling changes, targeted knock out of GPR37L1 or transient expression of certain rare variants produced alterations in cellular cholesterol homeostasis, suggesting a causal connection to the observed disease associations [19]. Finally, to simulate the impact of rare nonsense variants found in the large patient cohort, a knockout (KO) mouse line lacking *Gpr37L1* was generated, revealing loss of this receptor led to sex-specific behavioral changes implicated in migraine. Altogether, the human and mouse results focus attention GPR37L1 as a novel, astrocytic contributor [20] to migraine-related disorders.

## Methods

### Genomic analyses

To generate disease associations across the phenotype spectrum represented in the EHR, we used WES from 51,289 participants recruited through the MyCode® Community Health Initiative, which includes Geisinger patients enrolled through primary care and specialty outpatient clinics [21]. Participants, who consented to broad research use of genomic data in the DiscovEHR database linked to EHR entries, were 59% female, median age of 61 years, predominantly Caucasian (98%) and non-Hispanic/Latino (95%) [22]. Sample preparation, sequencing, sequence alignment, variant identification, genotype assignment and quality control steps were carried out as previously described [22,23]. Rare variants (MAF < 0.01) were sorted into nonsense and frame-shift variants (=LOF), and missense variants (=MISS). Missense variants were further triaged by bioinformatics (RMPath score [12]) to predict benign, likely benign, likely pathogenic and pathogenic variants, which were binned for subsequent analyses. Details of the bioinformatics triage are as previously reported [12]. De-identified EHR data obtained from an approved data broker was sorted to identify all unique ICD-9 codes with ≥200 patients having at least 3 independent incidences of the code [12]. Individuals having no calls of each code were set as controls. Non-Europeans and one individual from closely related pairs up to first cousins were excluded from analyses. Models were adjusted for sex, age, age^2^ and first 4 principal components. We used the sequence kernel association test with default weights (SKAT R package), to compare the burden of rare variants in cases and controls [12]. To define the directionality of the association, all individuals having rare variants in *GPR37L1* in 51,289 whole-exome sequences (WES) and having EHR codes (ICD9 or, ICD10) for migraine (346.*, G43.*) or epilepsy (345.*, G40.*) were identified and the RMPath scores of the variants determined, as previously described [12]. All variants were classified as ‘Variants of Unknown/Uncertain Significance” (VUS), i.e., benign (VUSB), likely benign (VUSLB), likely pathogenic (VUSLP) or pathogenic (VUSP), according to the net scores determined by the algorithm [12]. The primary focus of this study was migraine, but both migraine and epilepsy are diseases characterized by hyperexcitability [19], therefore genomic analysis was designed to assess the degree of variant overlap between the two phenotypes.

### Generation and culture of SK-N-MC *GPR37L1* knockout cell line

The SK-N-MC *GPR37L1* knockout cell line was generated by the CRISPR/Cas9 method [24] The gRNA was designed using the Optimized CRISPR Design online tool (http://crispr.mit.edu). One high scored gRNA was found in *GPR37L1* exon one, 5’-GCCCTAACCCCGGCAAGGAT GGG-3’. This gRNA was cloned into pSpCas9(BB)-2A-Puro (PX459) V2.0, (Addgene, #62988). The positive clone was amplified to get the plasmid after sequence analysis. The SK-N-MC cell line (ATCC, #HTB-10^TM^) was cultured in Eagle’s Minimal Essential Medium (EMEM) Medium (ATCC) with 10% fetal bovine serum (FBS) (Gibco). On the day of electroporation, cells were detached by 5 min treatment with Stempro Accutase (Gibco) and isolated by centrifugation. 5x10⁶ cells were suspended in 100 µl of electroporation buffer with 20 µg plasmid, electroporated (Neon™ electroporation system, manufacturer’s suggested conditions), and seeded on 3-10 cm cell culture dishes (Corning). After 24 hours, puromycin (Sigma-Aldrich, 1 μg/ml) selection was initiated for 48 hours. Puromycin-resistant colonies were picked mechanically using sterile 20 µl tips and each colony was maintained in a separated well of a 48 well plate. *GPR37L1* knockout clone(s) were identified by PCR and sequence analysis (forward primer, 5’ GCCCTAACCCCGGCAAGGAT-3’; reverse prime, 5’-AGTAGCTGTGCCACACGATGCACATGAC-3’). Thirteen KO clones were identified by sequence analysis. The knockout clones were screened for off-target effects by Sanger sequencing for suspected genic sites that were identified by the Optimized CRISPR Design online tool. *In-silico* prediction determined the resemblance of genomic regions to the gRNA with up to 3 nucleotide mismatch tolerance; in total, three coding sites were analyzed. No alternate target modifications were observed in any of the GPR37L1-null clones compared to the sequence of the original SK-N-MC clone. Clone #40 was identified without off-target effects and was chosen for use in experimental studies (*GPR37L1*-KO SK cells). This clone has a 94bp deletion starting 3’ of the CRISPR-Cas9 cleavage site. The deletion caused a DNA coding sequence shift, resulting in a premature stop codon (TGA) in the sequence.

### Isolation and culture of primary mouse astrocytes

Primary mouse astrocytes were cultured from postnatal day 1-2 C57BL6 pups. Mice were quickly decapitated, and brains were removed and submerged in ice-cold dissection medium containing 1 mM sodium pyruvate, 0.1% glucose in HEPES-buffered saline, pH=7.4. The meninges were removed, and cortices were isolated and finely minced. A single-cell suspension was obtained by enzymatic digestion with 0.25% trypsin at 37°C for 20 min followed by trituration. Cells were seeded in a T75 flask in Dulbecco’s Minimal Essential Medium (DMEM) culture medium supplemented with 10% FBS and 1% penicillin-streptomycin antibiotics (Invitrogen). The medium was changed twice weekly. When cells became confluent, microglia and oligodendrocytes were removed by shaking at 260 rpm for 3 hours at 37 °C in an orbital shaker. Prior to biochemical assays, astrocytes were sub-cultured into different plates as indicated.

### Generation of WT and *GPR37L1* variants

The human *GPR37L1* cDNA clone (Origene) was subcloned into pcDNA3.1+ (Invitrogen) and used as the template to insert the Glu-Glu epitope sequence (EYMPME) and the GPR37L1 variants used in the experiments by *in vitro* mutagenesis using Q5 Site-Directed Mutagenesis Kit (BioLabs). cDNA clones were confirmed by Sanger DNA sequencing (GeneWiz). Primer sequences are available upon request.

### Western Blotting

WT GPR37L1 and variants (without or with the Glu-Glu tag) were transfected into *GPR37L1*-KO SK cells (SK-N-MC Transfection Kit, Altogen Biosystems). Cells were lysed after 48 hours, adjusted for equivalent amounts of protein using the BCA protein assay (Thermo Scientific), then 200 μg of lysate was incubated with anti-Glu-Glu tag antibody (Thermo Scientific) overnight at 4°C. The next day, the immunoprecipitated complexes were pulled down by protein G beads and eluted with 5x SDS-sample buffer. Eluted samples were run on 10% TGX gels (BioRad), transferred to PVDF membrane (BioRad), blocked with 5% milk in TBS-T, and exposed overnight at 4°C to anti-GPR37L1 polyclonal antibody (Abcam #151518). Blots were exposed to secondary HRP-conjugated antibody (GE Healthcare), developed with SuperSignal West Pico Chemiluminescence Substrate (Thermo Scientific), and imaged on an Odyssey Fe Imager (LiCor) and processed with Image Studio. When untagged or natively expressed receptors were studied by western blot, membrane preparations were prepared according to the Abcam Membrane Preparation Protocol, using 500 μL of cell fractionation buffer per 10 cm dish of confluent cells, scraped into glass tubes on ice (15 min). The cell suspension was passed 10x through a 25-gauge needle, incubated on ice (20 min), then particulates were pelleted in two rounds of centrifugation (3,000 rpm, 5 mins, 4°C), combining the resulting supernatants. Membranes were then pelleted from the combined supernatants by ultracentrifugation (85,000 rpm, 40 min, 4°C), and the pellet resuspended in buffer. Membranes were combined with 5x SDS-sample buffer and western blots performed as described above.

### Enzyme-linked Immuno-absorbance Assay (ELISA)

WT Glu-Glu tagged GPR37L1 and Glu-Glu tagged variants were transfected into *GPR37L1*-KO SK cells as described above. After 24 hours, equivalent numbers of cells were re-plated to poly-L-lysine-coated 96-well plates (Corning) and further cultured for 24 hours. In a subset of experiments, cells were treated for various times with 40 μM TX14(A) (Tocris). Cells were fixed with either MeOH or 4% paraformaldehyde for 15 minutes on ice; all further steps were carried out at room temperature. Cells were rinsed with TBS-T, blocked in 1% milk/TBS-T (1 hr), then incubated with anti-Glu-Glu tag antibody (1 hr), washed, then incubated with HRP-conjugated antibodies (1 hr). Samples were developed with TMB Liquid Substrate supersensitive solution (Sigma) for 30 minute and stopped with 1M sulfuric acid. Absorbances were read at 450 nm on ClarioStar (BMG Labtech). Nontransfected cells were used to determine background and blanks and background were subtracted from all samples. Three to four replicates for each condition were run in each experiment, which was repeated with at least three independent transfections.

### cAMP Assay

WT and variants were transfected into *GPR37L1*-KO SK cells. After 48 hours, cells were treated with 10 μM forskolin (Sigma) in DMSO and various concentrations (0-40 μM) of TX14(A) (Tocris), then cAMP levels were determined with the non-acetylated version of the direct cAMP ELISA kit (ENZO) following manufacturer’s guidelines. Absorbance signals at 405nm were read on a plate reader (ClarioStar, BMG Labtech). Optical densities were fitted to 4PL curve in Excel (Microsoft) and data analyzed with GraphPad PRISM (V.8.4.3). Raw data from three independent experiments run in triplicate were combined, averaged, and analyzed.

### Erk1/2 Phosphorylation Assay

GPR37-L1-KO SK cells was cultured in Eagle’s Minimal Essential Medium (EMEM) Medium (ATCC) with 10% fetal bovine serum (FBS) (Gibco). On the day of electroporation, cells were detached by 5 min treatment with Stempro Accutase (Gibco) and isolated by centrifugation. 5x10⁶ cells were suspended in 100 µl of buffer with 20 µg pcDNA3.1-GPR37L-1 plasmid, electroporated (Neon™ electroporation system, manufacturer’s suggested conditions), and seeded on 60mm cell culture dishes (Corning). After 24 hours, the cells were detached by 5 min treatment with Stempro Accutase and collected by centrifugation. The transfected cells were counted and 30,000 cells were seeded on each well in 96 well plate. The cells were incubated for another 24 hours, and then starved in 50 µl EMEM-Serum free medium (antibiotic free) in presence or absence of a protein kinase inhibitor (PKI) (6-22amide, 50 nM, GIBCO BRL) for 3 hours. The culture medium was removed out and the cells were washed with warmed PBS for 3 times and the PBS was removed out gently with pipette. Cells were treated with various concentrations (0-80 µM) of TX14(A) (TOCRIS) for 15 min in 50 µl of EMEM in presence or absence of PKI at 37 °C.

For the ERK1/2 phosphorylation assay, phospho- and total ERK1/2 were determined using the two-plate adherent cell protocol (Phosph-ERK1/2 kit, Cisbio), following manufacturer’s instructions. Fluorescent emission signals were read at two wavelengths (665 nm and 620 nm) on a plate reader (ClarioStar, BMG, Labtech), the gain of filter setting for 665 nm is at 2,000 and for 620 nm is at 1,700, and HTRF ratios were determined, and the resulting data was analyzed with GraphPad PRISM (V.8.4.3). Raw data from three independent experiments run in triplicate were combined, averaged, and analyzed.

### Total Cellular Cholesterol Assay

WT SK-N-MC cells, GPR37L1-KO cells, and GPR37L1-KO cells transiently transfected with WT or variant GPR37L1 cDNAs were split into 96-well plates and cultured overnight. Total cell cholesterol was measured by cleaving cholesterol esters with esterase then complexing for 30 minutes in the dark at 37°C with 50μl Amplex Red reagent/HRP/cholesterol oxidase working solution (Invitrogen). Fluorescence was measured on a plate reader (ClarioStar, BMG, Labtech). Values were fitted to an in-plate cholesterol standard curve. Each condition was run in quadruplicate in 3 independent experiments and raw data was averaged over all experiments.

## Generation and Validation of GPR37L1 knock-out mouse line

To simulate the impact of rare nonsense variants found in the large patient cohort, a knockout (KO) mouse line lacking *Gpr37L1* was generated using the CRISPR/Cas9 method [24]. The online tool (http://crispr.mit.edu) was used to identify a highly-scored sgRNA within exon 1 (5’-GGTACAGCACTATGTACCCGAGG-3’), which was subsequently cloned into the pSpCas9(BB)-2A-Puro (PX459) V2.0 vector (Addgene, #62988). After confirmation of positive clones by Sanger sequencing (GENEWIZ), the plasmid was injected into the pronuclei of one-cell-stage C57BL/6J embryos according to standard protocols and injected embryos were transferred to pseudopregnant mice (Transgenic and Gene Targeting Core, Georgia State University). From the eleven progeny, five mutant mouse pups were confirmed to have a single nucleotide deletion (C) at 3’ of the CRISPR-Cas9 cleavage site that introduced a premature stop codon (TGA). The *Gpr37L1^+/-^* male mice were mated with C57BL/6J female mice (Jackson Laboratory, Bar Harbor, ME) to produce mice carrying the deleted Gpr37L1 allele (Gpr37L1^-^), which were then intercrossed to produce the wild type, heterozygous, and homozygous knockout mice on a congenic background. To assess potential off target effects of CRISPR/Cas9 technology, the knockout mice were by direct Sanger sequencing for suspected sites that were identified by the Optimized CRISPR Design online tool. In-silico prediction determined the resemblance of genomic regions to the gRNA with up to 3 nucleotide mismatch tolerance; in total, three coding sites were analyzed and no alternate target modifications were observed in Gpr37L1 null mice.

For validation, genotyping was performed on genomic DNA obtained from tail clips, and PCR was performed using Taq DNA polymerase (New England Biolabs). Primers (forward primer 5’-ATCCATGCCAGACGCCTGAC); reverse primer (5’-ATCCCCGAGTAGCCTTTGCT) were used to amplify a 722bp fragment flanking the edited *Gpr37L1* sequence (Supplemental Figure 1). The amplified DNA fragment was digested with Ava1 (underlined) and subjected to agarose gel electrophoresis The PCR product from WT mice was digested into 2 fragments (322 and 400 bp) with Ava1, whereas the PCR products from the mutant mice could not be digested with Ava1. Subsequently, loss of the transcript and protein was confirmed by qPCR and immunoblot analysis on mouse brains from WT, heterozygote (het) and KO (hom) mice, using an anti-GPR37L1 polyclonal antibody (GeneTex, Cat. No is 100005).

### Animal Use

All animal experiments were performed in accordance with the National Institutes of Health Guidelines for the Care and Use of Laboratory Animals and were approved by the Institutional Animal Care and Use Committee of Florida Atlantic University. For behavioral studies, animals were bred on congenic C57BL/6J line, with *Gpr37L1* knockout mice designated as *Gpr37L1*^-/-^ and wildtype littermates designated as *Gpr37L1*^+/+^. All mice were group housed (3-4/cage) and used for experiments after they reached 9 weeks of age. Animals were kept on a 12-h light/dark cycle with lights off at 7:00 PM in a quiet temperature-controlled room (24°C-25°C).

### Drugs

CGRP (AnaSpec, Inc) was dissolved in PBS, used at the dose of 0.1 mg/kg [4; 5] and intraperitoneally (i.p.) administered in a 5 ml/kg volume injection.

### Periorbital mechanical allodynia

Behavioral assessment was conducted by an operator blinded to treatment conditions as we recently described [25,26]. In brief, following an acclimation period in which mice were gently handled, habituated to i.p. injection and held as during the test, two baseline pain measurements were conducted. To reduce variability only the second baseline was used for data analysis. Female and male mice were tested on separate days by the same operator. von Frey filaments of different forces (0.02, 0.04, 0.07, 0.16, 0.4, 1.0, and 1.4 g) were applied to the periorbital area perpendicular to the skin, with sufficient force to cause slight buckling, and held for approximately 2 sec to elicit a positive response. Mice were poked five times with the same filament in a uniform manner throughout the periorbital region before changing filament. The stimulation was initiated with the 0.16 g filament. A response occurred when the mouse stroked the face with its forelimb, withdrew its head from the stimulus or shook its head. Sensitivity was determined according to the up-down method. The 50% mechanical withdrawal threshold (expressed in g) was then calculated.

### Anxiety-like behavior

Anxiety-like behavior was assessed using the elevated plus maze (EPM) test as we described [25,26]. The EPM apparatus was a black, ‘plus’-shaped platform equipped with two open arms and two closed arms of the same dimensions (35.6 x 7.6 cm). The platform was at a raised height of 50 cm above the ground and placed under room lighting of 500 lux. The EPM test started when each mouse was placed on the central platform of the EPM apparatus faced to a closed arm and left free to explore the maze for 5 min. Each mouse performance was recorded by a micro camera set in the testing room and examined at a later time by an operator blinded to the treatment schedule. The operator recorded the time spent in the open arms of the maze as well as the entries into the open as measures of anxiety-like activity and the number of closed arm entries as an indicator of the mouse locomotor behavior.

### Statistical Analyses

Tibco Statistica (Version 13.5.0.17) was used for data analysis. Assessment of periorbital allodynia was analyzed as area under the curve (AUC) for the various treatment combinations followed by two-way ANOVA. Unpaired t test was used for analysis of anxiety-like behavior in the EPM. Significance was set at p<0.05.

## Results

### Disease associations of rare variants in *GPR37L1*

A computational approach was used to assess the clinical significance of *GPR37L1* genetic variants in 51,289 (51K) WES. Rare (*i.e.,* <u>M</u>ean <u>A</u>llele <u>F</u>requency (MAF) <0.001) *GPR37L1* variants were present in the heterozygous state and classified as loss-of-function (=LOF, nonsense or frameshift) and missense variants (=MISS) [12]. Although there were too few LOF variants for computational analysis, missense variants were categorized according to their predicted pathogenicity using a bioinformatics algorithm (i.e., RMPath) [12] and twenty-seven “likely pathogenic” or “pathogenic” variants were binned for further analysis. Subsequently, SKAT was performed by running the binned variants against all ICD9 codes with ≥200 individuals having three or more independent encounters of the code in their EHRs to ensure accuracy of disease diagnoses. As previously reported [12] and shown here (Table 1), this unbiased approach revealed the top disease association for *GPR37L1* variants was generalized convulsive epilepsy (ICD9 code 345.1; Q=6.29e-17), consistent with a published study linking *GPR37L1* with epilepsy in a consanguineous family [15]. Although the clinical phenotype of carriers was not reported, family members homozygous for the *GPR37L1* variant (p.Lys349Asn) developed a novel, progressive myoclonus epilepsy and early death by 20 years of age [12]. Also, pointing to the pleiotropic potential of this approach, epileptic seizures are commonly associated with traumatic mouth injuries [27] and *GPR37L1* variants were strongly linked to traumatic fracture of tooth (ICD9 code 873,63; Q=3.07e-12). Likewise, *GPR37L1* variants were associated with migraine (ICD9 code 346.1 migraine; Q=1.13e-8) and epilepsy and migraine are known to share many clinical features and pathological mechanisms [28]. Finally, attesting to the discovery promise of this approach, significant associations with malignant neoplasm of the kidney (ICD9 code 189) and unspecified disorder of kidney and ureter (ICD9 code 593.9) were also identified, underscoring the potential importance of GPR37L1 in vascular and renal function that has been suggested by mouse and human studies [29–33].

**Table 1.**
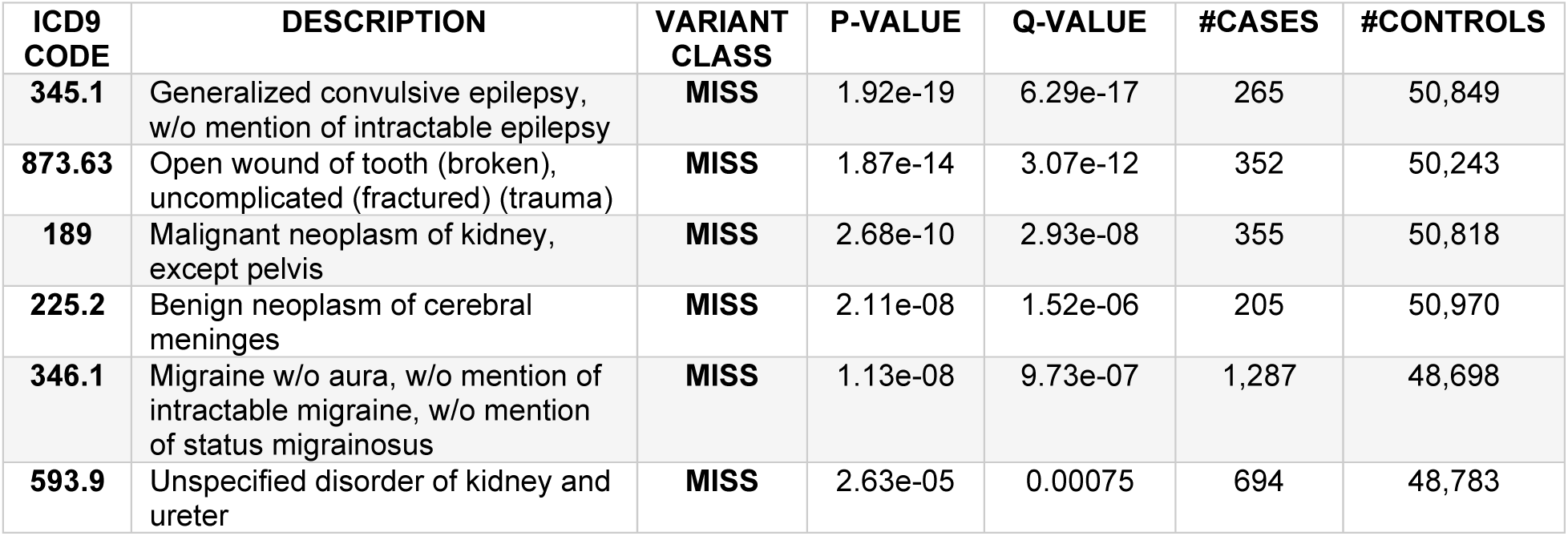
SKAT analysis of rare variants of *GPR37L1*. Missense (MISS) variants identified in 51,189 WES of the DiscovEHR cohort were binned and subjected to SKAT analysis against the set of ICD9 codes in the electronic health record (EHR) with at least 200 individuals having three or more independent instances of the code, regardless of genotype. Controls were specified as individuals without any instance of the code. Significance was defined as Q<e-5 with a stringent multiple-testing threshold of p=6.2e^-7^. The variants included in the analysis were: c.176A>G, p.Tyr59Cys; c.356C>T, p.Pro119Leu; c.362A>G, p.Tyr121Cys; c.365C>G, p.Pro122Arg; c.379T>G, p.Ser127Ala; c.388G>A, p.Ala130Thr; c.444C>A, p.Asn148Lys; c.449C>T, p.Ser150Leu; c.476A>G, p.Tyr159Cys; c.536T>C, p.Val179Ala; c.550C>T, p.Leu184Phe; c.560T>A, p.Val187Asp; c.571G>A, p.Glu191Lys; c.588G>T, p.Arg196Ser; c.598G>A, p.Asp200Asn; c668C>T, p.Ala223Val; c.680A>G, p.Asp227Gly; c.727G>A, p.Glu243Lys; c.730C>T, p.Arg244Trp; c.823G>A, p.Glu275Lys; c.886G>A, p.Glu296Lys; c.925C>G, p.Arg309Gly; c.1081G>A, p.Val361Met; c.1141G>A, p.Val381Met; c.1171C>T, p.Arg391Cys; c.1178C>T, p.Thr393Ile; c.1265C>T, p.Pro422Leu.

### Rare genomic variants in *GPR37L1* in individuals with migraine and/or epilepsy

The current study focused on addressing the impact of genetic variation within the coding sequence of *GPR37L1* on the incidence of migraine. Since migraine and epilepsy are both diseases of hyperexcitability and associated with *GPR37L1* rare variants, we performed clinical lookups of consented patients carrying *GPR37L1* variants with the relevant diagnoses in 51,289 WES, with the goal of assessing variant overlap and/or pleiotropy. First, we assessed the three individuals heterozygous for rare nonsense (LOF) variants and found that none had three validated calls of migraine or epilepsy in their EHRs. Two were males (age range 62-86) and one was female (age 77). Next, we captured all MISS variants and used the RMPath algorithm to predict their pathogenicity scores (Fig. 1A). Finally, we identified the smaller set of MISS variants found in individuals with at least 3 encounters for migraine (ICD9 code 346.*) or epilepsy (ICD9 code 345.*). Consistent with the relative prevalence of these disorders in the adult population [31], we found a larger number of MISS variants in individuals affected with migraine than epilepsy. Moreover, a smaller number of variants was identified in individuals with both diagnoses with significant overlap of variants but not in the same individuals, suggesting a common genetic basis for these conditions (Fig. 1A, overlap).

**Figure 1.**
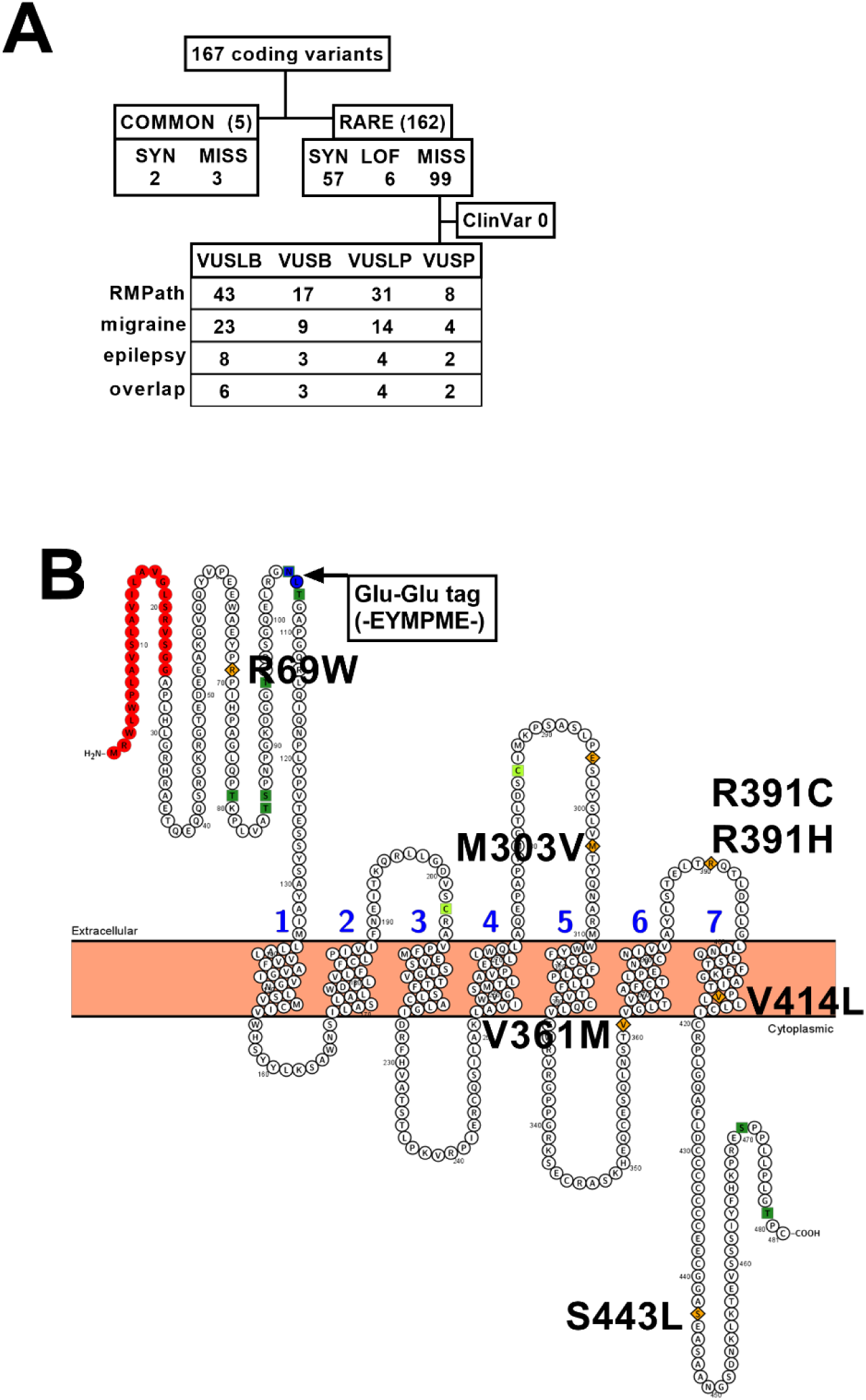
Pathogenicity scores of rare *GPR37L1* variants. **A.** Flow chart of *GPR37L1* rare variants in 51,289 WES, triaged by MAF, variant type (SYN=synonymous; LOF=loss-of-function nonsense or frameshift; MISS = missense). Bioinformatics triage was done by calculating RMPath scores [12] of all rare variants, and then identifying the subset of those found in migraine (346.*) and epilepsy (345.*) patients; range of scores was from -2 to 22; maximum possible score, 24. Scores were divided into 7 bins of equal width, B = benign, LB = likely benign, LP = likely pathogenic, and P = pathogenic. Overlap indicates rare variants found in both migraine and epilepsy patients (not necessarily in the same patient). **B.** Snake plot of GPR37L1 generated with Protter software [64] with locations of rare missense variants chosen for functional studies (p.Arg69Trp, p.Met303Val, p.Val361Met, p.Arg391His, p.Arg391Cys, p.Val414Leu, p.Ser443Leu). The location of the internal Glu-Glu tag is also indicated. Red = signal peptide residues; orange = variants studied; light green = cysteines participating in extracellular disulfide bond.

Since both migraine and epilepsy are complex disorders with genetic and environmental contributors [28], the penetrance of one or the other condition in individuals carrying *GPR37L1* variants is difficult to estimate. From the EHRs, we extracted both sex and age at first encounter for migraine and epilepsy. As shown in Table 2, carriers with a migraine diagnosis showed a significant bias toward females (∼79%), while those with an epilepsy diagnosis exhibited comparable incidence among males and females (61%). The DiscovEHR 51,289 WES dataset is slightly biased toward females, i.e., 59% female [22]. Thus, the relative number of variants and the differential sex bias of the two phenotypes reflect the distribution of these disorders in the adult population and examination of carriers with migraine illustrates that sex is an additional contributor.

**Table 2.** Demographics of individuals in 51,289 WES dataset with *GPR37L1* variants. All individuals with rare *GPR37L1* missense variants with specific ICD9 codes were identified and their sex and age of onset (first incidence of ICD9 code in the EHR) of the disorder. Only parental ICD9 codes are presented.

| **ICD9**<br>**code** | **description** | **#variants** | **#PTs** | **%Female** | **Age @ first**<br>**code** |
| --- | --- | --- | --- | --- | --- |
| 345* | epilepsy | 17 | 28 | 61% | 0-76 |
| 346* | migraine | 50 | 114 | 79% | 13-79 |

### SK-N-MC cells as a viable model system to study GPR37L1 expression and function

*GPR37L1* is highly expressed in astrocytes, and to a lesser extent, in neurons, microglia, and oligodendrocyte precursors [17,18]. Accordingly, we identified neuroblastoma-derived SK-N-MC cells, which endogenously express this receptor, as a viable cell model to assess the impact of genetic variation on GPR37L1 expression and function. SK-N-MC cells (hereafter abbreviated SK cells) are easily transfected and used broadly for studies of cellular signaling. To provide a null background, the CRISPR/Cas9 method was used to engineer SK cells lacking GPR37L1 (KO), as confirmed by PCR screening (Figure 2A) and immunoblotting analyses (Figure 2B). To assess GPR37L1 signaling in SK cells, we tested fragments of the multi-domain secreted protein prosaposin (PSAP), which has been identified as the endogenous ligand of GPR37L1 in some studies [16,18], but contradicted in others [35,36]. The PSAP protein is composed of four structurally similar modules, termed saposin A-D, and saposin C (SapC) is implicated in activation of GPR37L1. Because GPR37L1 is G_i/o_ coupled [18], we determined the efficacies of SapC domain, or a synthetic peptide fragment derived thereof, TX14(A) [16] on inhibition of forskolin-stimulated cAMP production. GPR37L1-KO SK cells transfected with WT GPR37L1 were stimulated with forskolin (FSK, 10 μM) in the absence or presence of varying concentrations of either SapC or TX14(A). As shown in Figure 2C, both the endogenous and synthetic agonists inhibited FSK-activated cAMP generation with similar IC_50_s (12.9 and 7.1 μM, respectively) although SapC produced a more robust inhibition of cAMP generation. TX14(A)-mediated inhibition of cAMP production is present in WT SK cells but notably absent in *GPR37L1*-KO SK cells (Figure 2D). Likewise, TX14(A) showed a greater than 50% reduction in FSK-mediated cAMP production in primary astrocytes (Figure 2E). Taken together, these results demonstrate that WT SK cells express GPR37L1 as the sole target of TX14(A) and thus *GPR37L1*-KO SK cells provide the appropriate cellular context for comparison of signaling by WT and variant GPR37L1.

**Figure 2.**
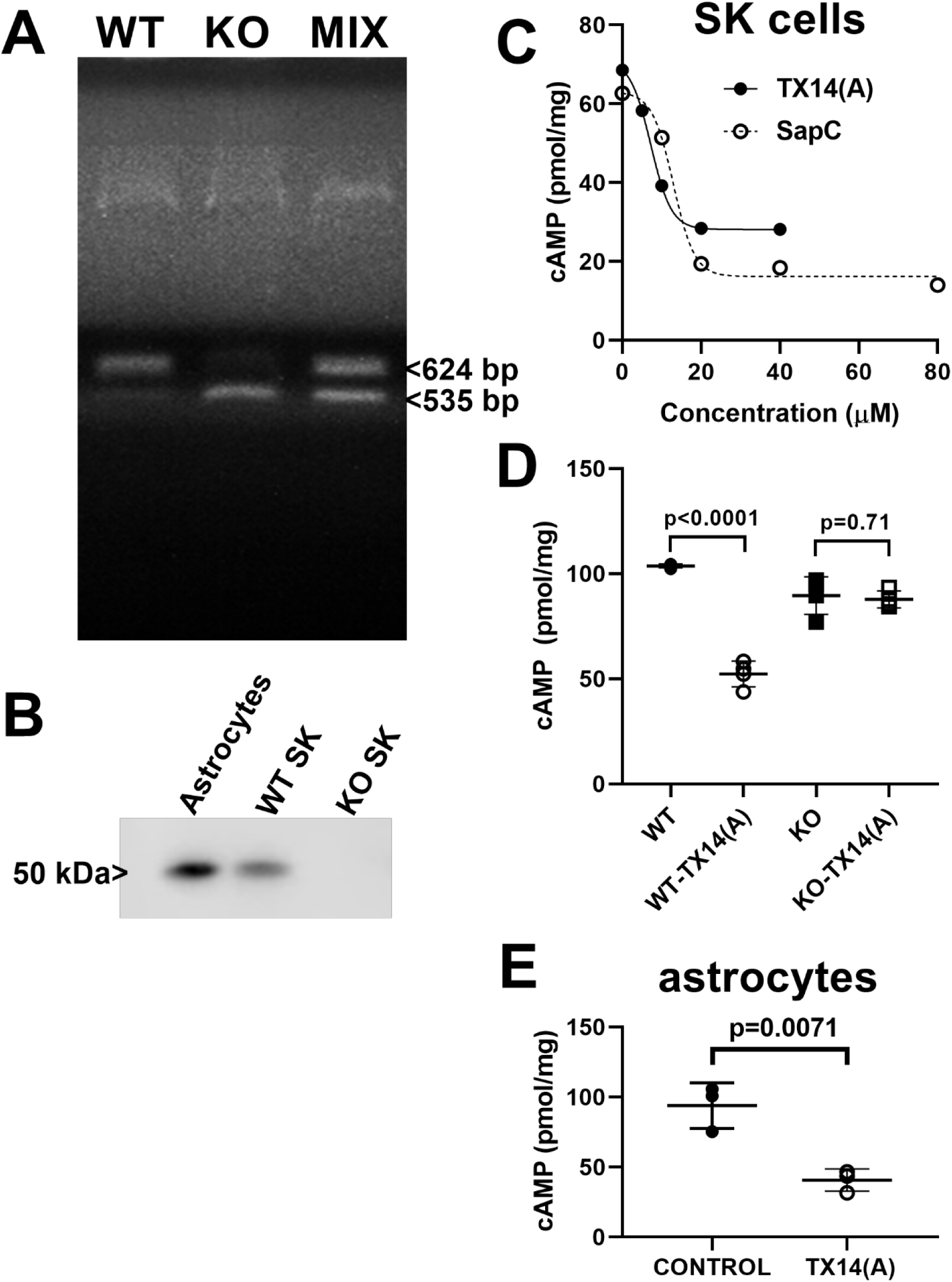
GPR37L1 expression and function in SK-N-MC cells. **A.** RT-PCR demonstrating endogenous expression of GPR37L1 in SK-N-MC cells (WT), and RT-PCR validation of a CRISPR/Cas9-generated *GPR37L1*-KO SK cell clone (KO). Final lane shows gel run of mixture of both PCR products (MIX). **B.** Western blot of membrane preparations from primary mouse astrocytes, WT and *GPR37L1*-KO SK cells, probed with anti-GPR37L1 antibody (Abcam). Membranes were isolated as described in Methods; 30 μg of membranes per lane. **C.** The PSAP module SapC and TX14(A) (Thr-D-Ala-Leu-Asp-Asn-Asn-Ala-Thr-Glu-Glu-Ile-Leu-Tyr) [16]) activate GPR37L1. Control SK-N-MC cells were stimulated with 10 μM forskolin ± different concentrations of the peptide TX14(A) (IC_50_ 7.2 μM) or SapC (IC_50_ 12.9 μM). cAMP was analyzed as described in Methods. Curves were generated by fitting all raw data from 3 independent experiment. **D.** WT and *GPR37L1*-KO cells were stimulated with 10 μM forskolin ± 40 μM TX14(A) and cAMP levels were assayed as described in Methods. **E.** Primary mouse astrocytes were stimulated with 10 μM forskolin ± 40 μM TX14(A) and cAMP levels were assayed as described in Methods. Significances in D & E were determined by two-tailed t-test assuming unequal variances, * p<0.05, ** p<0.01, *** p<0.001, **** p<0.0001.

### Functional analysis of GPR37L1 variants associated with migraine

For functional analyses, we chose GPR37L1 variants across the range of predicted pathogenicity located at sites conserved in human and mouse to broaden the long-term impact of these studies, i.e., VUSB, p.Met303Val and p.Val414Leu, VUSLB, p.Arg69Trp, p.Ser443Leu and p.Arg391His and VUSLP, p.Val361Met and p.Arg391Cys. Fortuitously, two variants with different predicted pathogenicity were identified at position 391, i.e., p.Arg391His (likely benign, RMPath score 8) in a single individual having no instances of relevant phenotypes, and p.Arg391Cys (likely pathogenic, RMPath score 15), found in 2 individuals, 1 with migraine. Table 3 summarizes the variants chosen and patient demographics, if any. Since bioinformatic predictions cannot account for variant-mediated changes of secondary or tertiary structure or at binding sites, the variant locations are marked in relation to the topology of the receptor (Figure 1B), i.e., in the amino terminal domain (p.Arg69Trp), the e2 loop (p.Met303Val), at the junction of the i3 loop with helix 6 (p.Val361Met), the e3 loop (p.Arg391His, p.Arg391Cys), at the cytoplasmic side of helix 7 (p.Val414Leu), and in the carboxyl terminal segment (p.Ser443Leu).

**Table 3.** *GPR37L1* migraine-associated variants selected for functional studies. Indicated are the number of homozygous or heterozygous individuals (HO_HE) in 51,289 WES with the variant (rs ID indicated) and details of the individuals with the indicated calls for epilepsy (345.*) and/or migraine (346.*). **<u>BOLD</u>** text indicates an individual with both phenotypes. Two variants had no individuals with indicated phenotypes and were included to test the range of pathogenicity scoring on function.

| VARIANT<br>rs ID | RMPath<br>class | RMPath<br>score | HO_HE | 345.* | 346.* |
| --- | --- | --- | --- | --- | --- |
| p.Arg69Trp<br>rs1219056911 | VUSLB | 9 | 0_5 | **F/21yrs** | **F/20yrs** |
|  |  |  |  |  | F/22yrs |
| p.Met303Val<br>rs771579982 | VUSB | 15 | 0_4 | none | F/23yrs; F/23yrs |
| p.Val361Met<br>rs114687119 | VUSLP | 11 | 0_6 | none | None |
| p.Arg391Cys<br>rs772132535 | VUSLP | 15 | 0_2 | none | F/37yrs |
| p.Arg391His<br>rs773384673 | VUSLB | 8 | 0_1 | none | None |
| p.Val414Leu<br>rs141962107 | VUSB | 4 | 0_6 | none | F/27yrs; F/15yrs |
| p.Ser443Leu<br>rs748607219 | VUSLB | 7 | 0_3 | none | None |

Functional analyses of these variants were performed in KO SK cells expressing either WT or variant receptors and the relative abilities of TX14(A) to attenuate FSK-stimulated cAMP production [16] were measured (Figure 3). We noted four distinct changes in signaling. Two receptor variants (p.Met303Val (VUSB), p.Arg391His (VUSLB), had signaling properties indistinguishable from the WT receptor in the presence or absence of TX14(A) (Figure 3Aa). Two other receptor variants (p.Val414Leu (VUSB), p.Ser443Leu (VUSLB)) were more effective at reducing cAMP than WT receptor in the presence of agonist (Figure 3Ab). Likewise, two receptor variants (p.Val361Met (VUSLP), p.Arg391Cys (VUSLP)) were more effective at reducing cAMP than WT receptor in the presence of TX14(A), but also showed significantly reduced cAMP in the absence of agonist, indicative of basal constitutive activity (Figure 3Ac). The final receptor variant (p.Arg69Trp (VUSLB)) was less effective at inhibiting cAMP than WT receptor in the presence of TX14(A) (Figure 3Ad). Figure 3B summarizes the diverse range of basal and TX14(A)-inhibited cAMP levels as well as apparent alterations relative to WT for each variant. Overall, these results show only a weak correlation between bioinformatics-based predictions of pathogenicity and experimental analysis of receptor variants, with analysis of receptor activity revealing altered signaling properties of most receptor variants.

**Figure 3.**
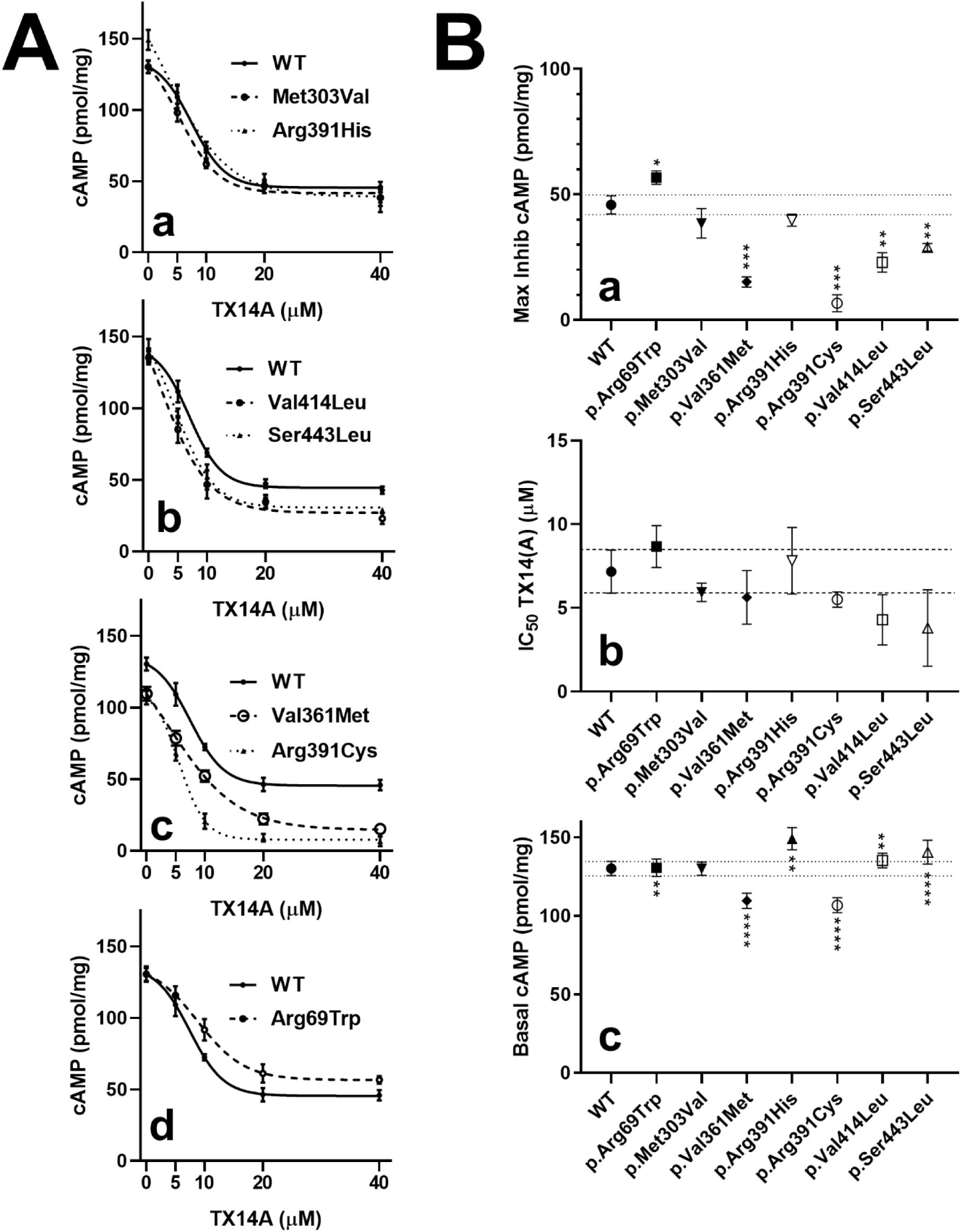
Migraine-associated variants in transfected *GPR37L1*-KO cells have differential ability to inhibit cAMP production. **A**. Migraine-associated variants were assessed for their ability to attenuate FSK-stimulated cAMP production when activated by TX14(A). *GPR37L1*-KO cells were transiently transfected with WT or variants, and net cellular cAMP was determined at a range of TX14(A) concentrations in cells stimulated with 10 μM forskolin, as described in Methods. Plots present averaged raw data ± SD results of three independent transfections. Variants are plotted with WT (solid line in all graphs), according to their different patterns of response. **a.** p.Met303Val (dashed line), p.Arg391His (dotted line). **b.** p.Val414Leu (dashed line), p.Ser443Leu (dotted line). **c.** p.Val361Met (dashed line), p.Arg391Cys (dotted line). **d.** p.Arg69Trp (dashed line). **B.** Quantitation of at least 3 independent experiments as illustrated in **A**. Inhibition curves were fitted and the maximal inhibition by TX14(A) (**a**), the IC_50_ for TX14(A) (**b**) and basal cAMP levels ((**c**), zero TX14(A)), were determined as described in Methods. Statistical differences were determined by two-tailed Student’s t-test, compared with WT responses, with * p<0.05, ** p<0.01, *** p<0.001, **** p<0.0001.

GPR37L1 also regulates mitogen-activated protein kinase (MAPK) pathways [16], therefore we next tested the ability of WT and receptor variants to activate ERK1/2 phosphorylation (Figure 4A). We noted three patterns of signaling alterations. Three receptor variants, two with normal cAMP inhibition (p.Met303Val, p.Arg391His) and one with increased potency for cAMP inhibition (p.Ser443Leu) had agonist-induced ERK1/2 phosphorylation comparable to WT receptor (Figure 4Aa). Three receptor variants (p.Val414Leu, p.Val361Met, p.Arg391Cys) with enhanced cAMP inhibition were found to have reduced ERK1/2 phosphorylation relative to WT in response to agonist (Figure 4ab). The last variant (p.Arg69Trp), with lower cAMP inhibition had an apparent increase in V_max_ (Figure 4Ac) which was not well-fitted over the accessible TX14(A) concentration range, although there was a significant, reproducible increase in ERK1/2 phosphorylation at 60 and 80 μM TX14(A) (Figure 4Ac). Despite variable changes in V_max_, all receptor variants had fitted EC_50_s comparable to WT receptor. Finally, we tested whether there was interaction between the cAMP and MAPK pathways by measuring ERK1/2 phosphorylation in cells expressing WT GPR37L1 in the absence or presence of 40 and 80 μM protein kinase A inhibitor peptide (PKI). As illustrated in Figure 4C, addition of PKI had no impact on TX14(A)-induced ERK1/2 phosphorylation. Taken together, these results demonstrate that a subset of GPR37L1 variants found in patients with a migraine diagnosis showed changes in agonist-induced cAMP inhibition and ERK1/2 phosphorylation that may be indicative of signaling bias.

**Figure 4.**
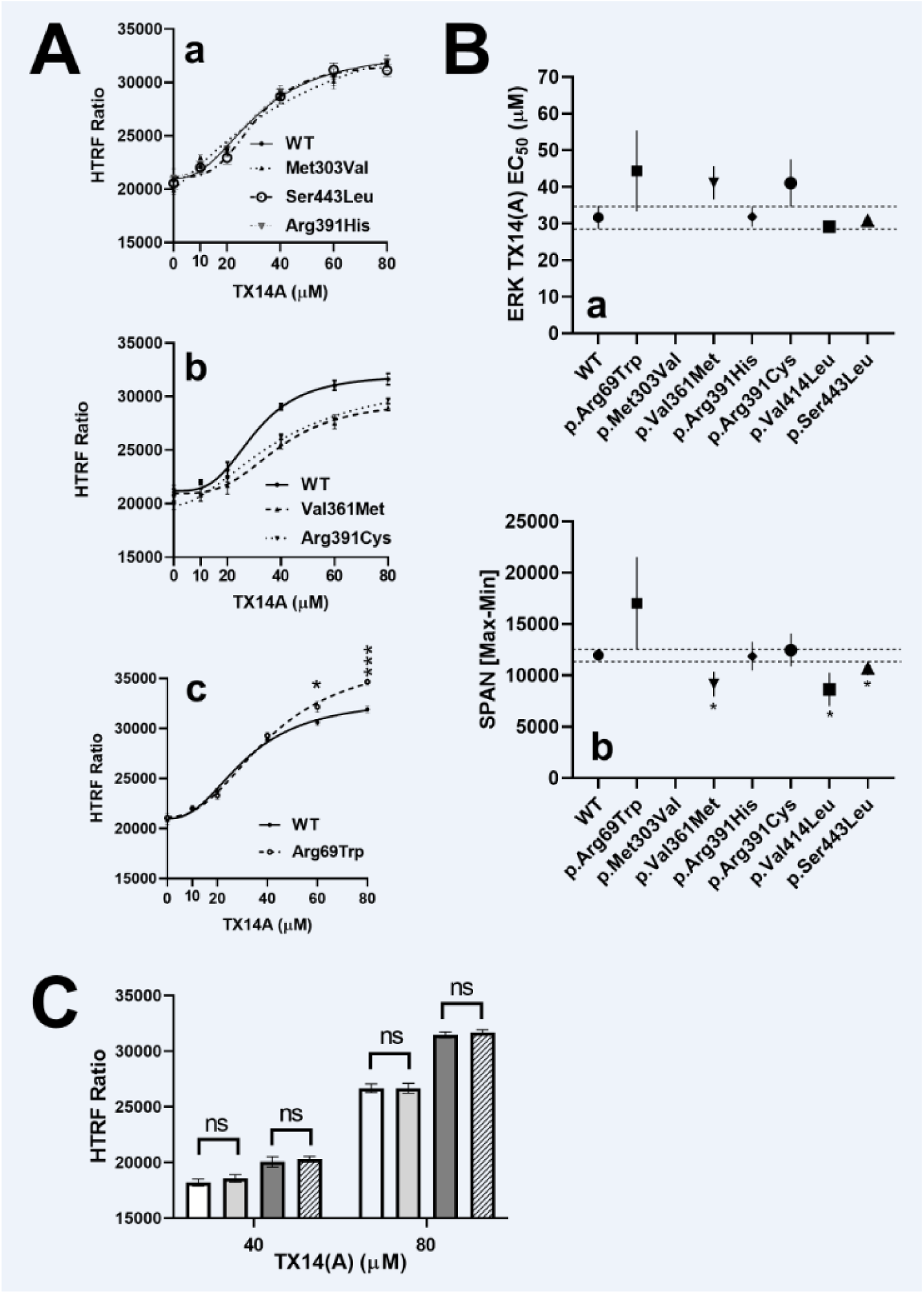
Migraine-associated variants in transfected *GPR37L1*-KO cells have differential ability stimulate ERK1/2 phosphorylation. **A.** The ability of TX14(A) to induce ERK1/2 phosphorylation in *GPR37L1*-KO cells transfected with WT or variants was determined in a plate-based assay as described in Methods. Plots illustrate mean of rad data ± SD of three independent transfections. Variants were plotted with WT (solid line) and sorted by their response differences. **a.** p.Met303Val (dashed line), p.Arg391His (dotted line), p.Ser443Leu (dash-dotted line). **b.** p.Val361Met (dashed line), p.Arg391Cys (dotted line), p.Val414Leu (dash-dotted line). **c.** p.Arg69Trp (dashed line). **B.** Stimulation curves were fitted and the EC_50_ estimates (**a**) and the span (maximum – minimum) (**b**) were determined as described in Methods. Statistical differences were determined by two-tailed Student’s t-test, compared with WT responses, * p<0.05, ** p<0.01, *** p<0.001, **** p<0.0001. **C**. Interaction of cAMP and MAPK pathways in GPR37L1 responses were tested by treatment with protein kinase inhibitor (PKI) showing its addition had no impact on TX14(A)-induced ERK1/2 as determined by one-way ANOVA analysis.

### Impact of migraine-associated variants on GPR37L1 expression and localization

To determine whether the signaling differences among GPR37L1 variants could in part be due to altered expression and/or subcellular distribution, we incorporated an epitope tag which allows net expression and plasma membrane targeting to be accurately assessed. Because GPR37L1 can be cleaved by endo-proteases at amino terminal sites [37,38], the epitope tag was inserted at a site in the amino terminus distal to potential GPR37L1 cleavage sites to ensure accurate assessment of expression levels. The small Glu-Glu epitope tag (-EYMPME) was introduced between residues p.Asn105 and p.Leu106 at the extracellular amino terminus (indicated in Figure 1B). Tagged WT GPR37L1 (=Glu-GPR37L1) allowed efficient immunoprecipitation (IP) of transfected receptors from KO SK cells. Figure 5A illustrates a western blot that compares the sizes of untagged WT GPR37L1 (MembWT) and WT Glu-GPR37L1 (IP). The blot was probed with a polyclonal anti-GPR37L1 antibody, and both Glu-GPR37L1 and the untagged WT GPR37L1 had a dominant band consistent with the predicted mass of the full-length receptor, with a minor band barely visible below. Glu-GPR37L1 had a slightly larger mass than the untagged version, as expected. Both WT Glu-GPR37L1 and the untagged WT GPR37L1 inhibited FSK-mediated accumulation of cAMP with comparable IC_50_s (Figure 5B). Thus, the Glu tag does not alter key features of the WT receptor and was therefore used to assess changes in expression and/or signaling by GPR37L1 variants.

**Figure 5.**
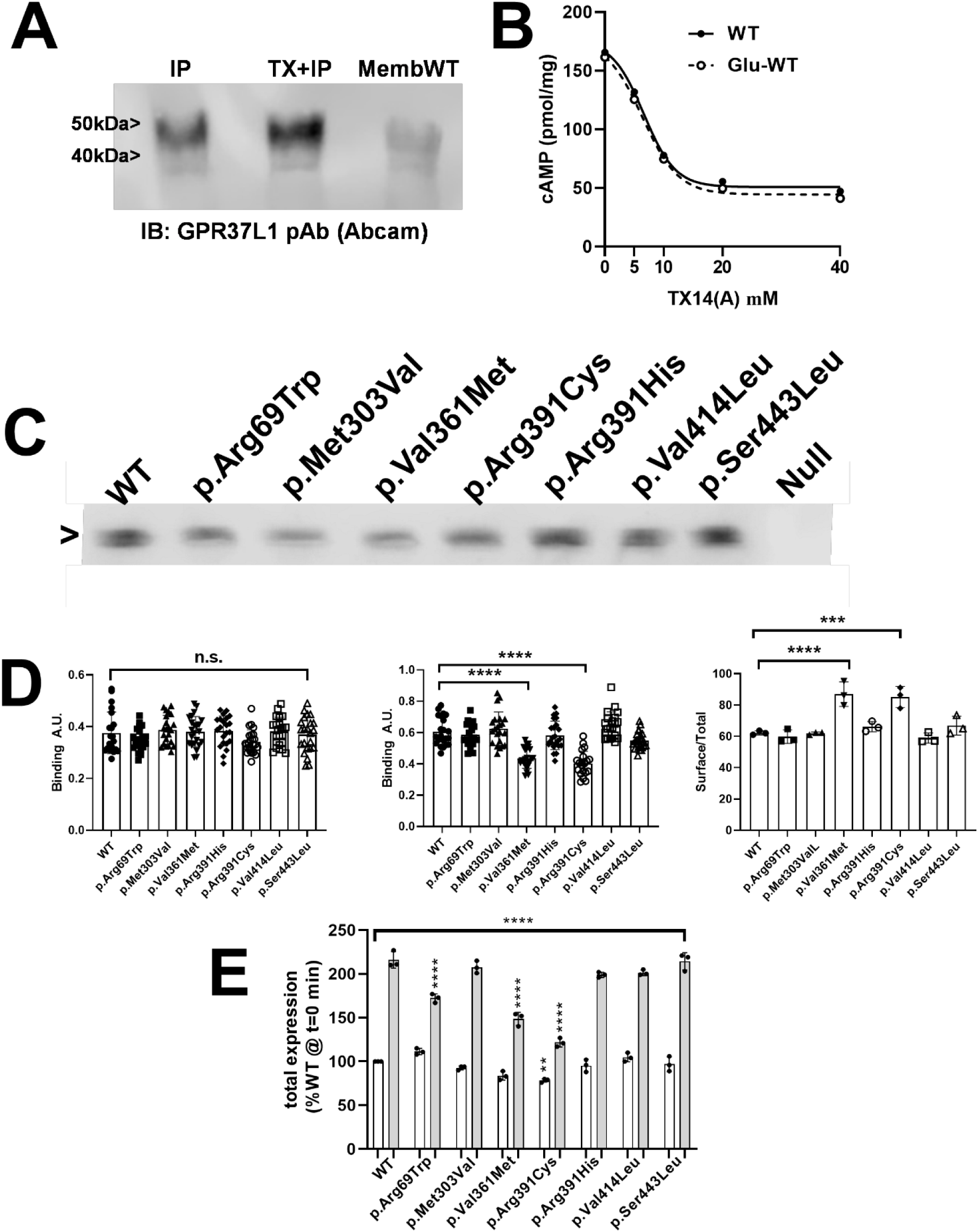
Variant-associated changes in GPR37L1 expression or localization. **A.** *GPR37L1*- KO SK cells were transfected with either untagged WT or Glu-GPR37L1, followed by immunoprecipitation with anti-Glu antibody or isolation of membranes to assess untagged GPR37L1. A second Glu-GPR37L1 transfected sample was treated ± 40 μM TX14(A) (30 min) prior to IP. Blot was probed with anti-GPR37L1 polyclonal antibody (Abcam). **B.** Response of GPR37L1-KO SK cells transfected with either WT or Glu-GPR37L1 to TX14(A) inhibition of FSK- mediated cAMP production. **C** GPR37L1-KO SK cells transfected with either WT or Glu-GPR37L1 to TX14(A) inhibition of FSK-mediated cAMP production. **C.** *GPR37L1*-KO SK cells transfected with Glu-tagged WT or variant GPR37L1, immunoprecipitated with anti-Glu antibodies and analyzed by western blot probed with anti-GPR37L1 antibody (see Methods). **D.** ELISA assays of Glu-tagged WT or variants assessing surface (left panel, fixed with paraformaldehyde) or total (middle panel, fixed with cold methanol) expression in the absence of TX14(A). Right panel illustrates the Surface/Total distribution of WT and variants. Data presents mean of all points (4 replicates x 3 independent experiments) or mean ± SD (right panel). Statistical analysis by two-way ANOVA with multiple comparisons, * p<0.05, ** p<0.01, *** p<0.001, **** p<0.0001. **E.** ELISA assays to assess total expression of GPR37L1 were used to compare basal (no TX14(A), white bars) and 30 min in 40 μM TX14(A) (gray bars) for WT and GPR37L1 variants. Three independent experiments were run in quadruplicate, and mean protein levels for each experiment were averaged and normalized to WT at zero time or after 30 min treatment with TX14(A). Data were analyzed by one-way ANOVA with multiple comparisons, and significance is indicated as * p<0.05, ** p<0.01, *** p<0.001, **** p<0.0001.

Detection of Glu-tagged WT and variant receptor was performed by western blotting (Figure 5C), while surface and total expression were quantified by ELISA (Figure 5D). Although most variants were comparably expressed to WT (surface expression, Figure 5Da; total expression, 5Db), two receptor variants (p.Val361Met and p.Arg391Cys) exhibited a reduced total expression relative to WT, and thus higher surface to total expression ratios (Figure 5Dc). As changes in surface expression of receptors can produce significant alterations in downstream signaling, it is interesting to note that both variant receptors also showed enhanced cAMP inhibition (recall Figure 3Ac). Surprisingly, we also found that short term treatment with TX14(A) produced an upregulation of GPR37L1 (Figure 5A). To explore the nature of this effect and to assess the potential impact of receptor variants, KO SK cells expressing Glu-tagged WT or variant receptors were treated with TX14(A) (30 min, 40 μM). Although the maximal extent varied, agonist treatment caused rapid, time-dependent increases in their expression, with receptor variants p.Val361Met and p.Arg391Cys having the weakest up-regulation (Figure 5E). It is of note that p.Arg391His has signaling equivalent to WT in both cAMP and ERK1/2∼P assays, and shows net expression and upregulation comparable to WT, while the p.Arg391Cys variant has enhanced basal activity and decreased EC_50_ in the cAMP inhibition assay, and reduced V_max_ in the ERK1/2∼P assay, and significantly less TX14(A)-mediated upregulation. Altogether, these results demonstrate the impact of GPR37L1 genetic variation on signaling efficacy, bias, and expression, with p.Arg391Cys and p.Val361Met showing changes across all three metrics. Results also reveal a rapid up-regulation of GPR37L1 expression in response to agonist treatment that is dependent upon receptor occupancy and/or signaling output(s).

### GPR37L1 and total cellular cholesterol levels

Perturbations in brain cholesterol homeostasis are linked to neurological disorders including migraine, in which hyperexcitability may play a role [19]. To assess the importance of GPR37L1 to cholesterol homeostasis, WT and KO SK cells, and KO cells transiently transfected with or without GPR37L1 receptor were used (Figure 6A). A comparison of total cholesterol content among these cell lines revealed that loss of endogenous expression of GPR37L1 significantly reduced cholesterol content (Figure 6B, compare WT and KO cells), while transient transfection of the WT receptor restored cholesterol abundance to a normal level. Of even more interest, acute treatment of cells expressing endogenous or exogenous GPR37L1 with the agonist TX14(A) for 30 min produced statistically significant increases in total cholesterol, which was not observed in KO cells lacking receptor (Figure 6B). These results point to a novel role for GPR37L1 in the acute regulation of cholesterol homeostasis in response to agonist.

**Figure 6.**
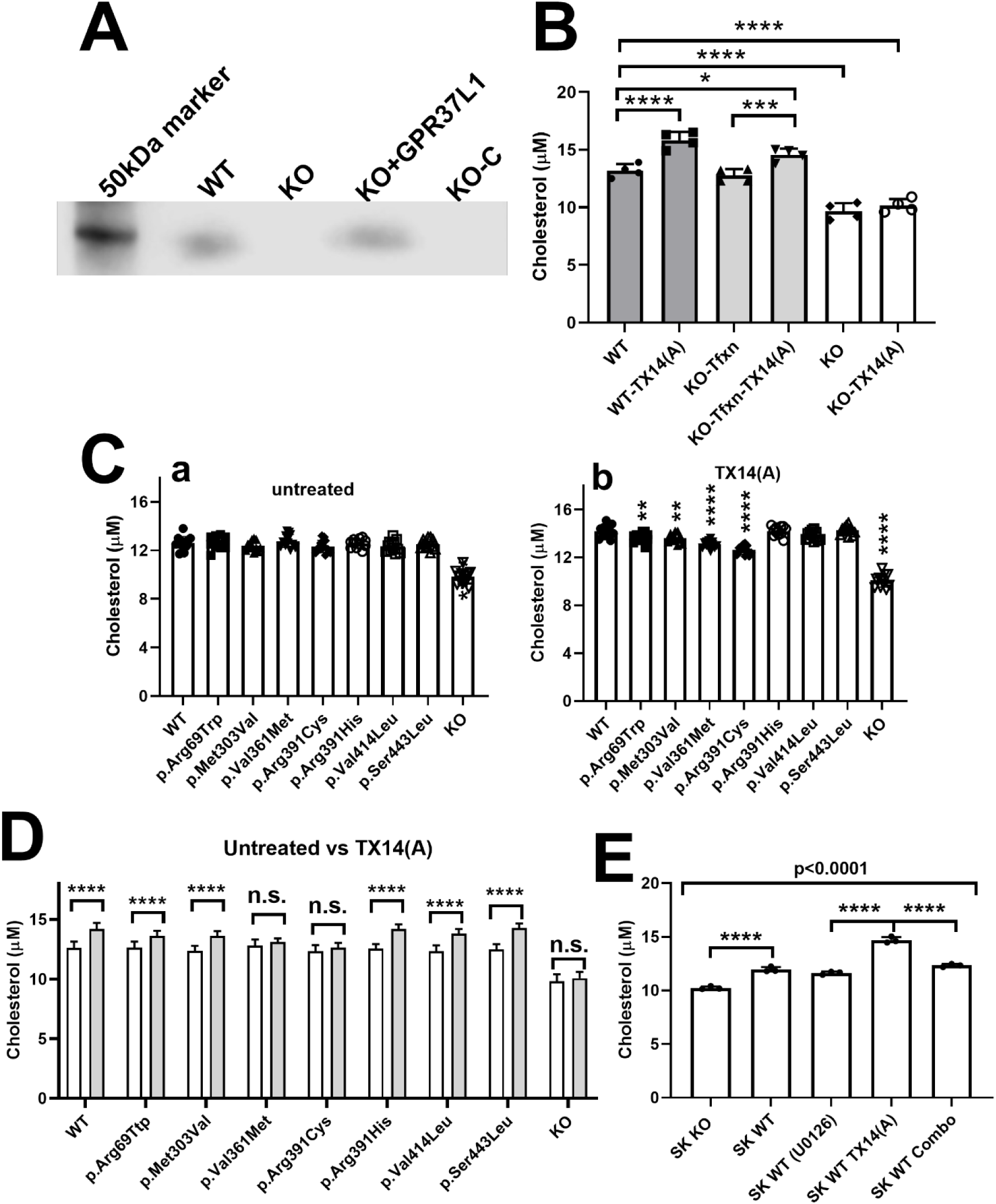
GPR37L1 regulation of cellular cholesterol levels. **A.** Western blot of WT SK cells, KO cells, KO cells transfected with WT GPR37L1, and KO cells exposed to transfection reagent (KO-C), blotted with anti-GPR37L1 antibody (Abcam). **B.** Total cellular cholesterol levels in WT SK cells, *GPR37L1*-KO cells transfected with WT GPR37L1, or *GPR37L1*-KO cells, treated with vehicle (DMSO) or 40 μM TX14(A) for 30 min. **C.** Total cellular cholesterol was measured in GPR37L1-KO cells transfected with WT or variants and treated with vehicle (**a**) or 40 μM TX14(A). (**b**) for 30 min. Analysis of **B. and C.** was performed by two-way ANOVA with multiple comparisons, relative to WT under the same condition, * p<0.05, ** p<0.01, *** p<0.001, **** p<0.0001. **D.** Comparison of total cellular cholesterol changes evoked by 40 μM TX14(A) treatment for 30 min, in WT or variant-transfected *GPR37L1*-KO cells. White, untreated, black, TX14(A)-treated. Analysis by Student’s t-test, * p<0.05, ** p<0.01, *** p<0.001, **** p<0.0001. E. Impact of U0126 on total cholesterol levels was assessed by treatment with either U0126 alone or in the presence of TX14(A) (=Combo), using WT SK cells. KO SK cells were used as a control, and treatment with U0126 had no effect on cholesterol levels in the absence of GPR37L1 (data not shown). Significance was determined by one way ANOVA with multiple comparisons, with * p<0.05, ** p<0.01, *** p<0.001, **** p<0.0001.

Since treatment with TX14(A) enhanced the differences in cellular cholesterol levels between WT and KO SK cells, we used the same approach to compare the total cellular cholesterol levels in KO cells transiently transfected with WT or variant GPR37L1. In the absence of TX14(A), cells expressing WT or receptor variants had equivalent amounts of total cell cholesterol, while there was an approximately 20% lower level in the KO SK cell line (Figure 6Ca). However, agonist treatment revealed differential responses between WT and receptor variants. Two receptor variants (p.Arg69Trp and p.Met303Val) showed weaker responses compared to WT, while two others (p.Val361Met and p.Arg391Cys) failed to increase total cholesterol levels in response to TX14(A) (Figure 6Cb). Variant responses to TX14(A) are plotted in Figure 6D. Of particular note, two receptor variants (p.Arg391Cys and p.Val361Met) with altered signaling properties in both cAMP and ERK1/2 assays failed to significantly increase total cholesterol levels in response to acute treatment with TX14(A). The different variants at position Cys391 again showed differential responses, i.e., TX14(A) elicited no cholesterol increase in cells expressing p.Arg391Cys while cells expressing p.Arg391His had an increase in total cholesterol in TX14(A) similar to WT.

Acute regulation of total cell cholesterol by GPCRs has not been fully characterized, although a recent study demonstrated NPY-mediated up-regulation of cholesterol in liver cells, a response that could be blocked by the MEK1/2 inhibitor U0126 [40]. Here, we determined whether the MAPK pathway contributed to TX14(A)-mediated up-regulation of total cholesterol in WT SK cells. As we have shown (Figure 4), TX14(A) increases ERK1/2 phosphorylation, and we hypothesized that blocking the upstream activator of ERK1/2 with U0126 should prevent TX14(A)-mediated increases in cellular cholesterol. Figure 6E illustrates that TX14(A) caused a significant increase in total cholesterol in WT SK cells. WT SK cells had higher total cholesterol levels than the KO SK cells, suggesting some constitutive activity of the endogenous GPR37L1 in WT SK cells. The U0126 had no effect on unstimulated WT or KO SK cells, but when treated with TX14(A), the U0126 severely attenuated the increase in cholesterol in WT SK cells. Overall, the results demonstrate that acute stimulation of GPR37L1 contributes significantly to total cellular cholesterol homeostasis through activation of MAPK pathways.

### Role of GPR37L1 in Migraine-Related Disorders

To simulate the impact of rare nonsense variants found in the large patient cohort, a knockout (KO) mouse line lacking *Gpr37L1* was generated on a congenic background to assess its role in neurological processes and permit dissection of the underlying mechanism(s) (Supplemental Figure 1). For the initial *in vivo* characterization, we focused on two migraine-related behaviors that share similar clinical features and suspected pathophysiology. For the acute migraine phenotype, we relied on a widely used model for migraine pathophysiology, acute CGRP injection, which produces facial pain sensitivity as measured by von Frey monofilaments [25]. In this case, KO and WT littermates showed increased pain sensitivity following a single CGRP injection that was indistinguishable between the two genotypes in female and male mice. Two-way ANOVA, conducted to analyze possible differences in allodynic responses calculated as AUC in GPR37L1^-/-^ vs. GPR37L1^+/+^ female mice treated with or without CGRP, led to a significant main effect of CGRP treatment [F(1,46)=4.4, p=0.04] accompanied by lack of genotype effect [F(1,46)=0.6, p=0.38] and “treatment x genotype” interaction [F(1,46)=0.0, p=0.76] (Figure 7A). In male mice, there was a significant effect of CGRP treatment [F(1,45)=18.3, p=0.0001], which, similarly to the outcome in females, was not accompanied by significant genotype effect [F(1,45)=1.0, p=0.315] or interaction [F(1,45)=0.4, p=0.534] (Figure 7B). Collectively, these results suggest that a single CGRP treatment evoked periorbital mechanical allodynia in male and female mice. However, the loss of Gpr37L1 did not affect the response to acute CGRP treatment.

**Figure 7.**
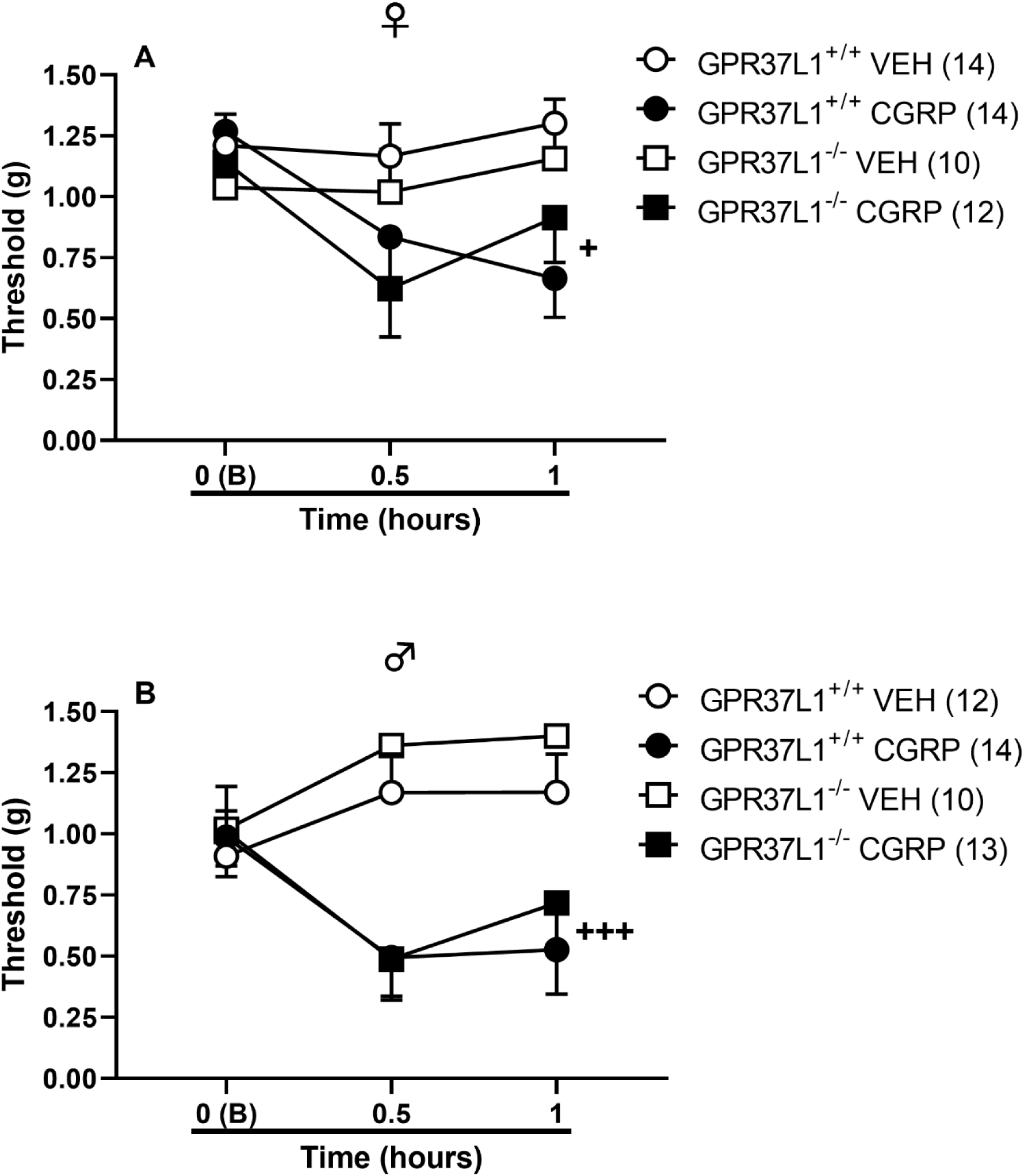
GPR37L1 knockout (GPR37L1^-/-^) mice and their wildtype littermates (GPR37L1^+/+^) show similar responses in an experimental model of migraine. Following acclimation, **(A)** female and **(B)** male GPR37L1^-/-^ and GPR37L1^+/+^ mice were tested for two basal thresholds of periorbital mechanical allodynia using the up-down method and then i.p. injected with calcitonin gene-related peptide (CGRP, 0.1 mg/kg) or VEH. In the CGRP model, both GPR37L1^-/-^ and GPR37L1^+/+^ female as well as male mice showed reduced thresholds of periorbital mechanical allodynia following CGRP treatment. Data are the mean sensitivity thresholds (grams) ± SEM 10-14 (total mice: 99) mice per group. *p<0.05 difference from GPR37L1^+/+^ groups; +p<0.05, +++p<0.001 difference from VEH-treated groups.

For the anxiety-related characterization, we used an established paradigm, the elevated plus maze (EPM), which measures time spent in the open vs enclosed arms [25,26]. To assess the effect of genetic disruption of the GPR37L1 receptor on anxiety phenotype, we compared their behavior in the EPM paradigm. Unpaired t test revealed that female GPR37L1^-/-^ mice significantly spent less time than WT mice on the open arms of the EPM t(20)=4.9, p<0.0001 (Figure 8A). In contrast, male GPR37L1^-/-^ mice showed similar open arm time to WT (GPR37L1^+/+^) mice t(14)=0.9, p=0.359 (Figure 8B). Mutant GPR37L1^-/-^ female mice also entered the open arms of the maze less than GPR37L1^+/+^ mice t(20)=3.7, p=0.0014 (Figure 8C), whereas males GPR37L1^-/-^ mice performed similarly to WT t(14)=1.2, p=0.238 (Figure 8D). GPR37L1^-/-^ mice of both sexes exhibited similar exploration of the maze, as shown by similar entries onto the closed arm of the EPM (females: t(20)=0.4, p=0.685; males: t(14)=2.0, p=0.0613, Figure 8E & 8F). Altogether, these results suggest that GPR37L1 deletion leads to sexually dimorphic anxiety-like responses, with female GPR37L1^-/-^ mice exhibiting increased anxiety-like activity as compared to WT controls.

**Figure 8.**
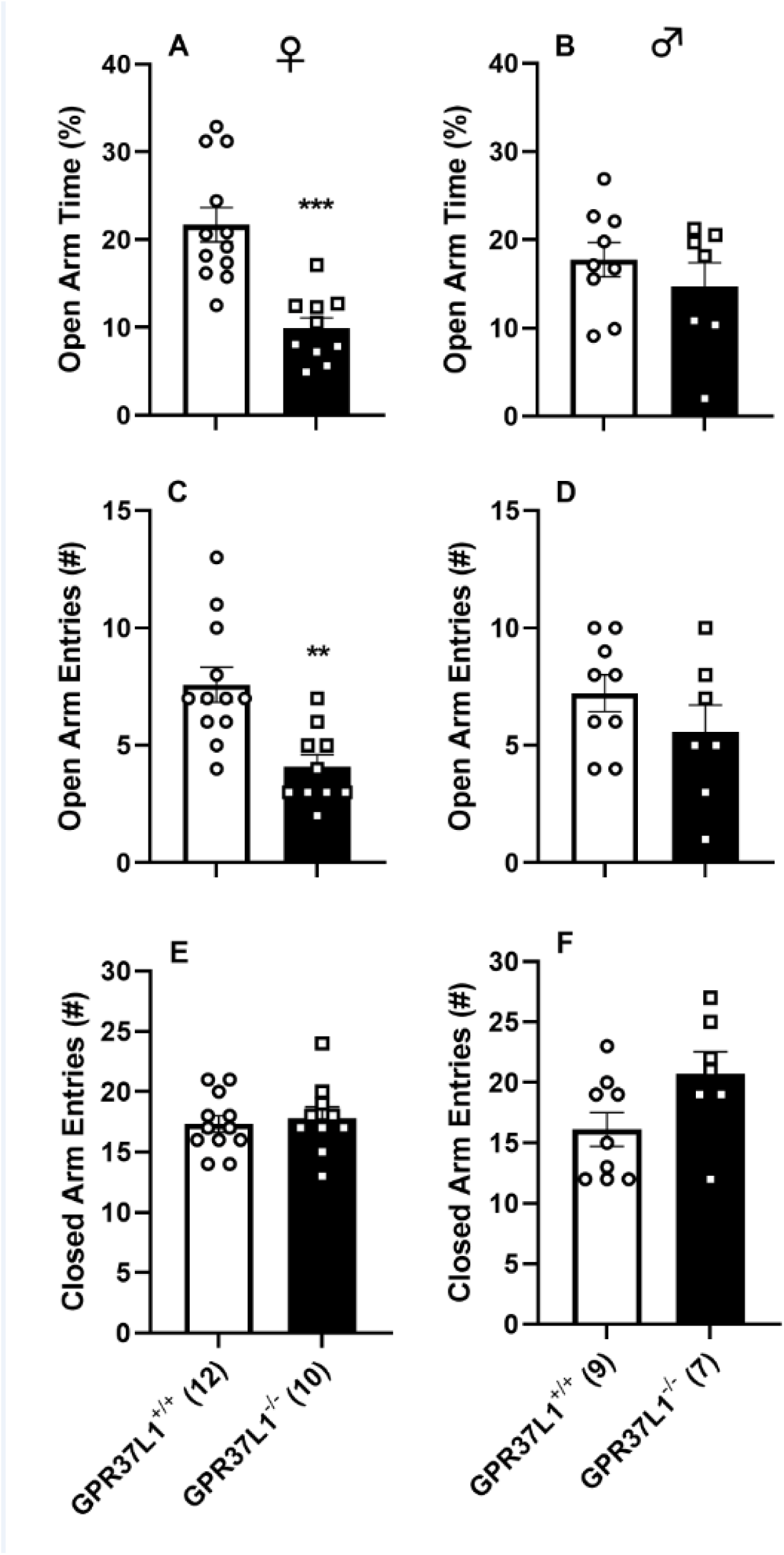
Sex differences in anxiety-like behavior for GPR37L1^-/-^ mice. **(A)** Female but not **(B)** male GPR37L1^-/-^ mice spent less time than GPR37L1^+/+^ mice exploring the open arms of the elevated plus maze (EPM), a behavior consistent with increased anxiety. Similarly, **(C)** female but not **(D)** male GPR37L1^-/-^ mice showed a reduced number of entries onto the open arms of the maze than GPR37L1^+/+^ mice, a second measure of anxiety-like behavior. **(E&F)** For both sexes, differences in anxiety parameters were accompanied by a similar number of closed arm entries (similar exploration of the EPM maze) for the two phenotypes. Values are presented as mean percent (%) open arm time and mean number of open and closed arm entries ± SEM of 10-12 (female) and 7-9 (male) GPR37L1^-/-^ vs. GPR37L1^+/+^ mice. **p<0.01, *** p<0.001 difference between phenotypes.

## Discussion

### Identifying GPR37L1 as a contributor to migraine and epilepsy risk

Using an unbiased computational approach, i.e. SKAT, we found that binned GPR37L1 coding variants were significantly associated with both epilepsy and migraine diagnoses. This was particularly interesting given the growing recognition that both disorders are strongly comorbid and may share common pathogenic mechanisms [28,34,41,42]. Although previously linked to a familial form of epilepsy [15], the linkage to migraine in the general patient population was both novel and highly significant for its strong sex bias. The latter may reflect the higher frequency of this diagnosis in the female portion in the DiscovEHR cohort and the general population in general [1,2], but additional studies will be needed to parse out the potential interaction(s) between sex and GPR37L1 function. Since SKAT can detect disease associations driven by only a subset of the binned coding variants, we then performed bioinformatics and functional analyses to identify variant receptors with altered functional properties. Notably, we identified some variant receptors whose properties were indistinguishable from wild type receptor. However, other variant receptors had significantly impacted properties, including those with missense mutations in critical domains, i.e., p.Arg69Trp in the extended amino terminal domain, p.Val361Met at the intracellular junction of the e3 loop and helix 6, and p.Arg391Cys in the e3 loop. Altogether, these results highlight both the need and value for performing integrated computational, bioinformatics, and functional analyses to reveal novel players and pathways for common diseases with multigenic inheritance [43–45].

### Validation of PSAP as the endogenous ligand of GPR37L1

GPR37L1 has been deorphanized as the receptor for secreted prosaposin, a neuroprotective protein that has a complex role in lysosomal function, being either directly transferred to the lysosome or targeted there after secretion by interactions with manose-6-phosphate receptors [47]. In the lysosome, prosaposin is cleaved into saposins A-D, with distinct roles in regulation of lipid metabolism [48,49]. Mutations in prosaposin cause lysosomal storage diseases, including Gaucher’s Disease from defects in SapC [50]. There is, however, controversy in the assignment of prosaposin as the endogenous agonist of GPR37L1, arising from studies primarily in heterologous expression systems, where GPR37L1 has been suggested to be constitutively active, and not regulated by the peptide agonist, TX14(A), which comprises the activating sequence identified in SapC [35–37]. Subsequent studies using KO mouse models clearly demonstrated, however, that loss of GPR37L1 eliminated the signaling responses elicited by TX14(A) [47]. Our studies in SK cells, confirm that WT SK cells express GPR37L1 and respond to both TX14(A) and the native SapC module, *GPR37L1*-KO SK cells do not respond to TX14(A), and transfection of *GPR37L1*-KO-SK cells with WT GPR37L1 restores the ability of TX14(A) to inhibit cAMP production. We demonstrate that the SapC module of PSAP or TX14(A), the peptidomimetic of SapC, activate GPR37L1, results consistent with PSAP being the endogenous agonist of GPR37L1.

### Dissection of functional differences among GPR37L1 variants

Functional studies of GPR37L1 variants were done in the null background of *GPR37L1*-KO SK cells, which possess all the regulatory and signaling partners for GPR37L1 function, as the WT cells express endogenous GPR37L1. We used standard functional assays, i.e., inhibition of FSK-induced cAMP production and stimulation of ERK1/2 phosphorylation, previously identified as signaling outputs of GPR37L1 [16]. We found a differential effect of the rare variants on the two pathways, suggesting that a subset of variants alter signaling bias, which has been observed for variants in other GPCRs. Table 4 summarizes the signaling changes identified for the GPR37L1 variants. Of note, short (15-30 min) exposures to TX14(A) induced up-regulation of GPR37L1 expression, a property observed for a subset of GPCRs involved in feed-forward regulation, including the Calcium Sensing Receptor [51,52]. Integrating this insight into models of astrocyte regulation will be critical to understanding the importance of GPR37L1 in the underlying pathologies of migraine and epilepsy.

**Table 4.**
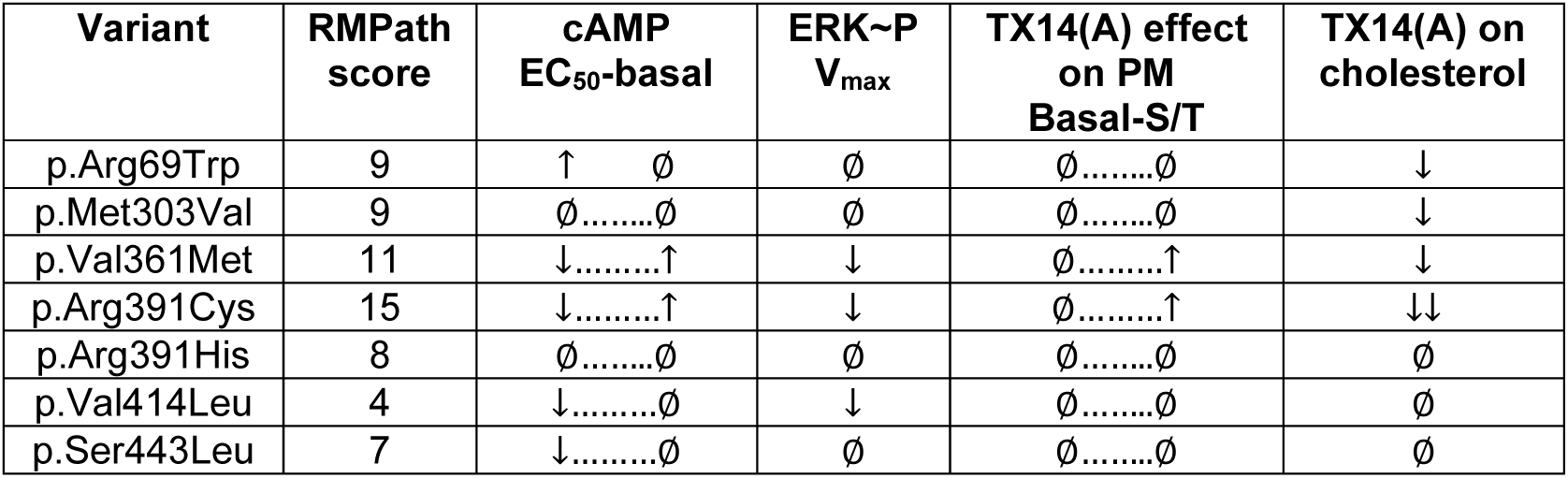
Summary of functional differences between WT and GPR37L1 variants. Variant functional results for TX14(A)-mediated inhibition of FSK-mediated cAMP production (EC_50_ for TX14(A) and change in basal FSK-mediated cAMP production in the absence of TX14(A)), TX14(A)-mediated ERK1/2∼P (fitted V_max_), the TX14(A)-mediated increase in expression of GPR37L1 (basal indicates level of expression in absence of TX14(A), S/T indicates the surface/total expression ratio after 30 min in 40 μM TX14(A)), and TX14(A) impact on cholesterol (the ability of 40 μM TX14(A) (30 min) to increase total cellular cholesterol levels).

### Identifying cellular role(s) for GPR37L1

The cellular role(s) for GPR37L in astrocytes is an area of active investigation. An agnostic RNAi screen for genes involved in cholesterol regulation identified PSAP as a regulator of perinuclear cholesterol levels in astrocytes [53]. Recent studies have shown that *Gpr37l1-KO* mice have altered cholesterol homeostasis in astrocytes, which contributes to dysregulation of Patched 1 internalization and trafficking [46]. In the current study, KO of GPR37L1 in SK cells decreased total cellular cholesterol by ∼20%, which was restored upon transfection with WT GPR37L1, suggesting GPR37L1 is a significant contributor to regulation of total cellular cholesterol. Whether native to the cells or re-expressed in KO SK cells, GPR37L1 increases cellular cholesterol, suggesting the possible presence of endogenous agonist. PSAP is widely expressed at low levels in most cell types, with low levels of secretion that can be upregulated after stress or injury [54]. Further, acute stimulation of WT or variant GPR37L1s with TX14(A) differentially up-regulated total cellular cholesterol, implicating GPR37L1 in the potential cholesterol crosstalk between astrocytes and neurons.

Up-regulation of cholesterol by acute stimulation of GPCRs has not been broadly studied at the cellular level, although a recent report suggests that neuropeptide Y stimulates cholesterol synthesis by acute activation of the SREBP2-HMGCR pathway in mouse hepatocytes [39]. Significant up-regulation of pathway proteins was observed within an hour, and proteins remained elevated for up to 24 hours [39]. Similar effects were observed in cultured hepatocytes, where pathway protein expression and cholesterol responses were sensitive to the MEK1/2 inhibitor U0126 [36]. Our results are consistent with MAPK regulation of acute cholesterol changes in SK cells, i.e., the ability of GPR37L1 variants to increase ERK1/2 phosphorylation is correlated with upregulation of cell cholesterol in response to TX14(A), and the MEK1/2 inhibitor U0126 significantly attenuates acute cholesterol up-regulation by TX14(A).

### Assessing *in vivo* role(s) for GPR37L1

Mouse models are critical in dissecting the mechanism(s) contributing to migraine. There are both common and rare forms of migraine, and human and mouse studies have begun to reveal the underlying mechanisms that account for the heterogeneity of this disorder [3–9]. One specific mechanism involves dysregulation of glutamate homeostasis[4,5]. While some rare monogenic disorders point to a direct role in neurons, other studies indicate a protective role in astrocytes to counteract the effects of such dysregulation [55,56]. Based on human clinical data, astrocyte localization, and functional studies, we posit a role for GPR37L1 signaling in the latter scenario. In mice, a single systemic administration of CGRP induces periorbital mechanical allodynia and other responses consistent with migraine including spontaneous pain, altered light sensitivity (photophobia) and anxiety-like behavior [25]. We examined here the impact of genetic disruption of the GPR37L1 receptor and found that such deletion did not affect the ability of an acute CGRP injection of eliciting acute cephalic allodynia. However, migraine is a complex disorder that extends beyond acute, episodic pain and involves functional and structural brain changes as well as increased responsiveness (sensitization) of central neurons responsible for pain, features that may underlie chronic migraine mechanisms [57,58].. Likewise, the impact of GPR37L1 may be increased in chronic migraine, as it has been noted that GPR37L1 is upregulated in microglia and astrocytes after nerve injury [59]. Consistently, dysfunction of brain astrocytes, cells that harbor the GPR37L1 receptor, has been shown to trigger migraine-like symptoms in mice [60]. Given the important role of GPR37L1 in astrocyte maturation and function [61], it is conceivable that the lack of GPR37L1 receptor may affect chronic migraine-like states more than episodic migraine evoked by acute challenge of migraine-provoking substances. In future experiments, we will examine the role of the GPR37L1 receptor in chronic migraine models leading to sensitization of pain responses [25].

In conclusion, the current studies began by using an unbiased approach to define the potential clinical disorders associated with GPR37L1 rare variants. Strong associations with migraine and epilepsy were identified, showing a significant sex bias for migraine incidence in females. In vitro studies of GPR37L1 variants identified signaling changes in multiple pathways with altered signaling bias, highlighting potential contributions to clinical phenotypes. Initial studies in a KO mouse model again demonstrate a sex bias for females in an anxiety-related trait, though no effect in an acute migraine model. These studies confirm the importance of GPR37L1 in episodic neurological disorders including anxiety and migraine, suggesting the potential for common underlying mechanisms driving these clinically important disorders.

## Acknowledgements

We thank Kirk Jeffreys for help with initial experiments, Dr. Diane Smelser for helpful discussions, and Geisinger for support in early stages of these studies including access to the MyCode clinical and genomic data. We acknowledge and appreciate the generosity of the Geisinger patients who agreed to allow their data to be used for research.

---

**Supplemental Figure 1.**
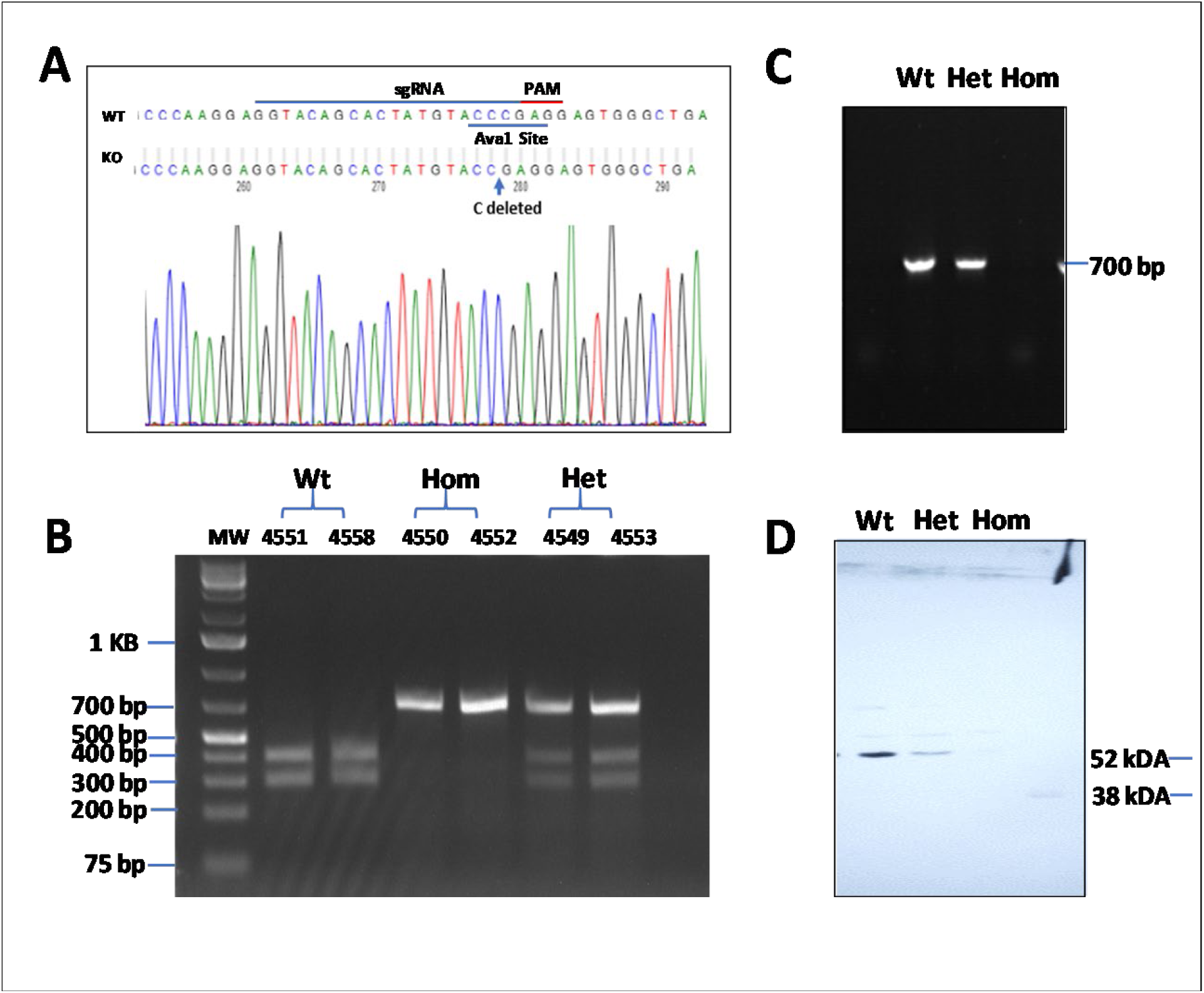
Generation and Validation of GPR37L1 KO mice. **(A)** Schematic representation of sgRNA targeting sequence and PAM site in the mouse GPR37L1 exon 1 region. The Ava1 recognition sequence is underlined, and a single base pair “C” deletion in GPR37L1 KO mice is indicated in sequence. **(B)** Genotyping of WT and GPR37L1KO mice. A 722 bp PCR product was amplified with a pair of primers that flanked the gene edited sequence of GPR37L1. The PCR product from WT mice was digested into 2 fragments (322 and 400 bp) with Ava1. However, the PCR products from the mutant mice could not be digested with Ava1. **(C)** RT-PCR shows the relative quantity of GPR37L1 mRNA extracted from mouse brains in the WT, heterozygote (het) and KO (hom) mice. **(D)** Western Blot analysis of protein expression in the brains of WT, het and KO mice.

## References

1. Burch, R.C., S. Loder, E. Loder, and T.A. Smitherman, The prevalence and burden of migraine and severe headache in the United States: updated statistics from government health surveillance studies. Headache, 2015. 55(1): 21–34.

2. Bonafede, M., S. Sapra, N. Shah, S. Tepper, K. Cappell, and P. Desai. Direct and indirect healthcare resource utilization and costs among migraine patients in the United States. Headache, 2018. 58(5): 700–714.

3. Sutherland, H.G., Albury, C.L., and L.R. Griffiths. Advances in genetics of migraine. J. Headache Pain. 2019. 20(1): 72.

4. Goadsby, P.J., Holland, P.R., Martins-Oliveira, M., Hoffmann, J., Schankin, C., and S. Akerman. Pathophysiology of migraine: a disorder of sensory processing. Physiol. Rev., 2017. 97(2): 553–622.

5. Noseda, R. and R. Burstein, Migraine pathophysiology: anatomy of the trigeminovascular pathway and associated neurological symptoms, CSD, sensitization and modulation of pain. Pain, 2013. 154 Suppl 1.

6. Mulder, E.J., C. Van Baal, D. Gaist, M. Kallela, J. Kaprio, et al., Genetic and environmental influences on migraine: a twin study across six countries. Twin Res, 2003. 6(5): 422–31.

7. Russell, M.B., L. Iselius, and J. Olesen, Inheritance of migraine investigated by complex segregation analysis. Hum Genet, 1995. 96(6): 726–30.

8. Gormley, P., V. Anttila, B.S. Winsvold, P. Palta, T. Esko, et al., Meta-analysis of 375,000 individuals identifies 38 susceptibility loci for migraine. Nat Genet, 2016. 48(8): 856–66.

9. Hansen, R.D., A.F. Christensen, and J. Olesen, Family studies to find rare high risk variants in migraine. J Headache Pain, 2017. 18(1): 32.

10. Tardiolo, G., P. Bramanti, and E. Mazzon, Migraine: Experimental Models and Novel Therapeutic Approaches. Int J Mol Sci, 2019. 20(12).

11. Manolio, T.A., F.S. Collins, N.J. Cox, D.B. Goldstein, L.A. Hindorff, et al., Finding the missing heritability of complex diseases. Nature. 2009. 461(7265): 747–53.

12. Dershem, R., Metpally, R.P.R., Jeffreys, K., Krishnamurthy, S., Smelser, D.T., Hershfinkel, M., Regeneron Genetics Center, Carey, D.J., Robishaw, J.D., and G.E. Breitwieser. Rare variant pathogenicity triage and inclusion of synonymous variants improves analysis of disease associations. J Biol Chem. 2019. 294(48): 18109–18121.

13. Tang, X.L., Y. Wang, D.L. Li, J. Luo, and M.Y. Liu, Orphan G protein-coupled receptors (GPCRs): biological functions and potential drug targets. Acta Pharmacol Sin. 2012. 33(3): 363–71.

14. Fricker, L.D. and L.A. Devi, Orphan neuropeptides and receptors: Novel therapeutic targets. Pharmacol Ther, 2018. 185: p. 26–33.

15. Giddens, M.M., J.C. Wong, J.P. Schroeder, E.G. Farrow, B.M. Smith, et al., GPR37L1 modulates seizure susceptibility: Evidence from mouse studies and analyses of a human GPR37L1 variant. Neurobiol Dis, 2017. 106: p. 181–190.

16. Meyer, R.C., Giddens, M.M., Schaefer, S.A., Hall, R.A. GPR37 and GPR37L1 are receptor for the neuroprotective and glioprotective factors prosaptide and prosaposin. Proc Natl Acad Sci USA, 2013. 110(23): p.9529–9523.

17. Jolly, S., Bazargani, N., Quiroga, A.C., Prongle, N.P., Attwell, D., et al. G protein-coupled receptor 37-like 1 modulates astrocyte glutamate transporters and neuronal NMDA receptors and in neuroprotective in ischemia. Glia, 2018. 66(1): 47–61.

18. Liu, B., Mosienko, V., Vaccari Cardoso, B., Prokudina, D., Huentelman, M., et al. Glio-and neuro-protection by prosaposin is mediated by orphan G-protein coupled receptors GPR37L1 and GPR37. Glia, 2918. 66(11): 2414–2426.

19. Hanin A., Baudin, P., Demeret, S., Roussel, D., Lecas, S., et al. Disturbances of brain cholesterol metabolism: A new excitotoxic process associated with status epilepticus. Neurobiol. Dis. 2021. 154: 105346.

20. Sutherland, H.G., Maksemous, N., Albury, C.L., Ibrahim, O., Smith, R.A., Lee, R.A., Haupt, L.M., Jenkins, B., Tsang, B., and L.R. Griffiths. Comprehensive exonic sequencing of hemiplagic migraine-related genes in a cohort of suspected probands identifies known and potential pathogenic variants. Cells, 2020. 9: 2368.

21. Carey, D.J., S.N. Fetterolf, F.D. Davis, W.A. Faucett, H.L. Kirchner, et al., The Geisinger MyCode community health initiative: an electronic health record-linked biobank for precision medicine research. Genet Med, 2016. 18(9): 906–13.

22. Dewey, F.E., Murray, M.F., Overton, J.D., Habegger, L., Leader, J.B., et al. Distribution and clinical impact of functional variants in 50,726 whole-exome sequences from the DiscovEHR cohort. Science, 2016. 354(6319): aaf6814. Doi:10.1126/science.aaf6814.

23. Abul-Husn, N.S., Manickam, K., Jones, L.K., Wright, E.A., Hartzel, D.N., et al. Genetic identification of familial hypercholersterolemia within a single U.S. health care system. Science, 2016. 354(6319):aaf7000. Doi:10.1126/science.aaf7000.

24. Mashiko D, Fujihara Y, Satouh Y, Miyata H, Isotani A, Ikawa M. Generation of mutant mice by pronuclear injection of circular plasmid expressing Cas9 and single guided RNA. Sci Rep. 2013 Nov 27;3:3355.

25. Preclinical effects of cannabidiol in an experimental model of migraine. Sturaro C, Fakhoury B, Targowska-Duda KM, Zribi G, Schoch J, Ruzza C, Calò G, Toll L, Cippitelli A. Pain. 2023 Jun 9. doi: 10.1097/j.pain.0000000000002960. Online ahead of print. PMID: 37310430

26. Targowska-Duda KM, Ozawa A, Bertels Z, Cippitelli A, Marcus JL, Mielke-Maday HK, Zribi G, Rainey AN, Kieffer BL, Pradhan AA, Toll L. NOP receptor agonist attenuates nitroglycerin-induced migraine-like symptoms in mice. Neuropharmacology 2020;170:108029.

27. Gawlak, D., Euniewska, J., Stojak, W., Hovhannisyan, A., Strozynska, A., et al. The prevalence of orodental trauma during epileptic seizures in terms of dental treatment – Survey study. Neurol. Neurochir. Pol. 2017. 51(5): 361–365.

28. Rogawski, M.A. Common pathophysiologic mechanisms in migraine and epilepsy. Arch. Neurol. 2008. 65(6): 709–714.

29. Mouat, M.A., Jackson, K.L., Coleman, J.L.J., Paterson, M.R., Graham, R.M. et al. Deletion of Orphan G Protein-coupled Receptor GPR37L1 in mice alters cardiovascular homeostasis in a sex-specific manner. Front. Pharmacol. 2021. 11:600266.

30. Mouat, M.A., Coleman, J.L.J., Wu, J., Dos Remedios, C.G., Feneley, M.P., et al. Involvement of GPR37L1 in murine blood pressure regulation and human cardiac disease pathophysiology. Am. J. Physiol. Heart Circ. Physiol. 2021. 321(4): H807–H817.

31. Zheng, X., Asico, L.D., Ma, X., and P.R. Konkalmatt. G protein-coupled receptor 37L1 regulates renal sodium transport and blood pressure. Am. J. Physiol. Renal Physiol. 2019. 316(3): F506–F516.

32. Coleman, J.L.J., Mouat, M.A., Wu, J., Jancovski, N., Bassi, J.K. et al. Orphan receptor GPR37L1 contributes to the sexual dimorphism of central cardiovascular control. Biol. Sex Differ. 2018. 9(1): 14.

33. Min, K.D., Asakura, M., Liao, Nakamura, K., Okazaki, et al. Identification of genes related to heart failure using global gene expression profiling of human failing myocardium. Biochem. Biophys. Res. Commun. 2010. 393(1): 55–60.

34. Huang, Y., Xiao, H., Qin, X., Nong, Y., Zou, D., Wu, Y. The genetic relationship between epilepsy and hemiplegic migraine. Neuropsychiatr. Dis. Treat. 2017. 13: 1175–1179.

35. Ngo, T., Wilkins, B.P., So, S.S., Keov, P., Chahal, K.K. et al. Orphan receptor GPR37L1 remains unliganded. Nat. Chem. Biol. 2021. 17(4): 383–386.

36. Coleman, J.L.J., Ngo, T., Schmidt, J., Mrad, N., Liew, C.K., et al. Metalloprotease cleavage of the N terminus of the orphan G protein-coupled receptor GPR37L1 reduces its constitutive activity. Sci. Signal. 2016. 9(423): ra36.

37. Coleman, J.L.J., Ngo, T., Smythe, R.E., Cleave, A.J., Jones, N.M., Graham, R.M., Smith, N.J. The N-terminus of GPR37L1 is proteolytically processed by matrix metalloproteases. Sci. Rep. 2020. 10:19995.

38. Mattila, S.O., Tuusa, J.T., Petaja-Repo, U.E. The Parkinson’s-disease-associated receptor GPR37 undergoes metalloprotease-mediated N-terminal cleavage and ectodomain shedding. J. Cell Sci. 2016. 129: 136601377.

39. Chen, F., Zhou, Y., Shen, M., Wang. Y. NPY stimulates cholesterol synthesis by activating the SREBP2-HMGCR pathway through the Y1 and Y5 receptors in murine hepatocytes. Life Sci. 2020. 262:118478.

40. Zarcone, D., Corbetta, S. Shared mechanisms of epilepsy, migraine and affective disorders. Neurol. Sci. 2017. 38(Suppl 1):73–76.

41. Shu, Y., Xu, Y., Xiao, W., Deng, X., Zeng, Y., et al. A conjoint analysis of epilepsy and migraine through network-and-pathway-based method. Ann. Palliat. Med. 2020. 9(5): 2642–2653.

42. Rogawski, M.A. Migraine and epilepsy-shared mechanisms within the family of episodic disorders. In: Noebels, J.L., Avoli, M., Rogawski, M.A., Olsen, R.W., Delgado-Escueta, A.V., editors. Jasper’s Basic Mechanisms of the Epilepsies (Internet). 4th edition, Bethesda (MD), National Center for Biotechnology Information (US); 2012.

43. Sun, H., and G. Yu. New insights into the pathogenicity of non-synonymous variants through multi-level analysis. Sci. Rep. 2019. 9:1667.

44. Ishikawa, T., Kimoto, H., Mishima, H., Yamagata, K., Ogata, S., et al. Functionally validating SCN5A variants allow interpretation of pathogenicity and prediction of lethal events in Brugada syndrome. Eur. Heart J. 2021. 42(29): 2854–2863.

45. Raraigh, K.S., Han, S.T., Davis, E., Evans, T.A., Pellicore, M.J., et al. Functional assays are essential for interpretation of missense variants associated with variable expressivity. Am. J. Hum. Genet. 2018. 102(6): 1062–1077.

46. La Sala, G., Di Pietro, C., Matteoni, R., Bolasco, G., Marazziti, G., and P. Tocchini-Valentini. Gpr37l1/prosaposin receptor regulates Ptch1 trafficking, Shh production, and cell proliferation in cerebellar primary astrocytes. J. Neurosci. Res. 2020. Doi: 10.1002/jnr.24775. PMID: 33350496.

47. Carvelli, L., Libin, Y., Morales, C.R. Prosaposin: a protein with differential sorting and multiple functions. Histol. Histopathol. 2015. 30(6): 647–660.

48. Leonova, T., Qi, X., Bencosme, A., Ponce, E., Sun, Y., Grabowski, G.A. Proteolytic processing patterns of prosaposin in insect and mammalian cells. J. Biol. Chem. 1996. 271(29): 17312–17320.

49. Hiraiwa, M., O’Brien, J.S., Kishimoto, Y., Galdzicka, M., Fluharty, A.L., Ginns, E.I., Martin, B.M. Isolation, characterization, and proteolysis of human prosaposin, the precursor of saposins (sphingolipid activator proteins). Arch. Biochem. Biophys. 1993. 304(1): 110–116.

50. Kang, L., Zhan, X., Ye, J., Han, L., Qiu, W., Gu, X., Zhang, H. A rare form of Gaucher disease resulting from saposin C deficiency. Blood Cells Mol. Dis. 2018. 68: 60–65.

51. Grant, M.P., Stepanchick, A., Cavanaugh, A., and G.E. Breitwieser. Agonist-driven maturation and plasma membrane insertion of calcium-sensing receptors dynamically controls signal amplitude. Sci. Signal. 2011. 4(200): fa78.

52. Breitwieser, G.E. Minireview: The intimate link between calcium sensing receptor trafficking and signaling: implications for disorders of calcium homeostasis. Mol. Endoccrinol. 2012. 26(9):1483–1495.

53 Bartz,F., Kern, L., Erz, D., Zhu, M., Gilbert, D., Meinhof, T., Wirkner, U., Erfle, H., Muckenthaler, M., Pepperkok, R., Runz, H. Identification of cholesterol-regulating genes by targeted RNAi screening. Cell Metab. 2009. 10:63–75.

54. GTEx Consortium. *Human genomics*. The Genotype-Tissue Expression (GTEx) pilot analysis: multitissue gene regulation in humans. Science. 2015. 348(6235): 648–660.

55. Capuani C, Melone M, Tottene A, Bragina L, Crivellaro G, Santellow M, Casari G, Conti F, Pietrobon D. Defective glutamate and K+ clearance by cortical astrocytes in familial hemiplegic migraine type 2. EMBO Mol Med. 2016; 8(8): 967–986.

56. Conti F, Pietrobon D. Astrocytic Glutamate Transporters and Migraine. Neurochem Res. 2023. 48(4):1167–1179.

57. Mungoven TJ, Henderson LA, Meylakh N. Chronic Migraine Pathophysiology and Treatment: A Review of Current Perspectives. Front. Pain Res. 2021 2: 1–15.

58. Brennan KC, Peitrobon D. A Systems Neuroscience Approach to Migraine. Neuron. 2018, 7(5): 1004–1021.

59. Kunihiro J, Nabeka H, Wakisaka H, Unuma K, Islam Khan S, et al. Prosaposin and its receptors GRP37 and GPR37L1 show increased immunoreactivity in the facial nucleus following facial nerve transection. PLoS One. 2020, 15(12): e0241315.

60. Romanos J, Benke D, Pietrobon D, Zeilhofer HU, Santello M. Astrocyte dysfunction increases cortical dendritic excitability and promotes cranial pain in familial migraine. Sci. Adv. 2020; 6: eaaz1584.

61. Nguyen T, Camp CR, Doan JK, Traynelis SF, Sloan SA, Hall RA. GPR37L1 controls maturation and organization of cortical astrocytes during development. Glia 2023; 71(8): C1, 1787-2066. https://doi.org/10.1002/glia.24375.

